# The impact of type VI secretion system, bacteriocins and antibiotics on competition amongst Soft-Rot *Enterobacteriaceae*: Regulation of carbapenem biosynthesis by iron and the transcriptional regulator Fur

**DOI:** 10.1101/497016

**Authors:** Divine Yutefar Shyntum, Ntombikayise Nkomo, Alessandro Rino Gricia, Ntwanano Luann Shigange, Daniel Bellieny-Rabelo, Lucy Novungayo Moleleki

**Affiliations:** Department of Biochemistry, Genetics and Microbiology, University of Pretoria, Lunnon Road, Pretoria, South Africa, 0028; Forestry, Agriculture and Biotechnology Institute, University of Pretoria, Lunnon Road, Pretoria, South Africa, 0028

## Abstract

Plant microbial communities’ complexity provide a rich model for investigation on biochemical and regulatory strategies involved in interbacterial competition. Within these niches, the soft rot *Enterobacteriaceae* (SRE) comprise an emerging group of plant-pathogens inflicting soft rot/black-leg diseases and causing economic losses worldwide in a variety of crops. In this report, a range of molecular and computational techniques are utilized to survey the contribution of antimicrobial factors such as bacteriocins, carbapenem antibiotic and type VI secretion system (T6SS) in interbacterial competition among plant-pathogens/endophytes using an aggressive SRE as a case study (*Pectobacterium carotovorum* subsp. *brasiliense* strain PBR1692 – *Pcb*1692). A preliminary screening using next-generation sequencing of 16S rRNA comparatively analysing healthy and diseased potato tubers, followed by *in vitro* competition assays, corroborated the aggressiveness of *Pcb*1692 against several relevant taxa sharing this niche ranging from Proteobacteria to *Firmicutes*. The results showed growth inhibition of several Proteobacteria by Pcb1692 depends either on carbapenem or pyocin production. Whereas for targeted *Firmicutes*, only pyocin seems to play a role in growth inhibition by *Pcb*1692. Further analyses elucidated that although T6SS confers no relevant advantage during *in vitro* competition, a significant attenuation in competition by the mutant strain lacking a functional T6SS was observed *in planta*. Furthermore, production of carbapenem by *Pcb*1692 was observably dependent on the presence of environmental iron and oxygen. Additionally, upon deletion of *fur, sly*A and *exp*I regulators, carbapenem production ceased, implying a complex regulatory mechanism involving these three genes. Potential Fur binding sites found upstream of *sly*A, *car*R and *exp*R in *Pectobacterium* genomes harboring carbapenem-associated genes further suggests a conserved regulatory pattern in the genus, in which carbapenem might be modulated in response to iron through the control exerted by Fur over secondary regulators. Furthermore, we unveiled the striking role played by S-pyocin in growth inhibition within the SRE group.

**Authors Summary:** For many phytopathogenic bacteria, more is known about interactions within the host and virulence factors used for host colonisation while relatively less is known about microbe-microbe interactions and factors that shape niche colonisation. The soft rot *Enterobacteriaceae* (SRE) comprise an emerging group of phytopathogens causing soft rot/black-leg diseases in a variety of crops leading to huge economic losses worldwide. In this report, a range of molecular and computational techniques are utilized to survey the contribution of antimicrobial factors such as bacteriocins, carbapenem antibiotic and type VI secretion system (T6SS) in interbacterial competition among plant-pathogens/endophytes using an aggressive SRE as a case study (*Pcb*1692). Our results show that *Pcb*1692 inhibits growth of other SRE and several potato endophytes using either the type VI secretion, carbapenem or bacteriocins. Carbapenem plays a role in both inter and intrabacterial competition *in vitro*, while the *Pcb*1692T6SS plays a role in interbacterial competition *in planta* (in potato tubers). We also demonstrate that carbapenem regulation requires the presence of environmental iron and oxygen in a complex network consisting of *Pcb*1692 Fur, SlyA, and ExpI. The presence of these gene homologs in several SREs suggests that they too can deploy similar antimicrobials to target other bacteria.

## Introduction

For phytopathogenic bacteria, more is known about interactions within their hosts and virulence factors that these microbes recruit to successfully colonise their hosts while relatively less is known about microbe-microbe interactions and how such interactions impact on niche colonisations (1-4). Studies on the mammalian gut microbiome, the advent of new sequencing technologies, availability of many ‘omics’ data sets such as metagenomics, transcriptomics and proteomics are some of the factors that have brought microbial interactions to the fore (5-9). Thus, the past few decades have seen a rapid growth in the body of literature on microbial interactions.

Bacteria exist in complex multispecies communities which are mostly characterised by competition and to a lesser extent, cooperation (10-12). In these interactions, survival depends on the ability to compete for resources in a given niche. Mechanisms of competition can be either classified as exploitative or interference (10, 13-16). Exploitative competition is passive whereby microbes compete for scarce resources and the winner typically limits nutrient availability from competitors. Interference competition is characterised by production of specialised metabolites. Microbes have a large arsenal of antimicrobial weapons that mediate interference competition and these include contact dependant inhibition (CDI), T6SS, antibiotic and bacteriocin production (17-29). These antimicrobials are diverse in their mechanism of action, targets, antagonistic and spatial range (20, 21). There is clearly a cost in the execution of these weapons, hence, by implication, regulation must be tightly modulated (26, 30-32). Over the years, many studies have interrogated mechanisms of the different antimicrobials ‘singularly’ however fewer have systematically investigated these factors in concert. For example a growing number of studies have shown the role, mechanism of action and targets of T6SS, T5SS and bacteriocins in bacterial competitions (22, 28).

Soft Rot *Enterobacteriaceae* (SRE), which were recently reclassified as Soft Rot *Pectobacteriaceae* (SRP), are a group of pathogens causing huge economic losses on potato production worldwide (33, 34). They cause blackleg and soft rot diseases in the field or during post-harvest storage, respectively. Decaying potato tubers are characterised by multispecies bacterial communities that are both Gram positive and negative (7, 33). Furthermore, within the SRE, it is quite normal to isolate bacteria from *Pectobacterium spp* as well as *Dickeya spp*, the two primary genera within *Pectobacteriaceae* from a single infected plant (7). Of these, *Pectobacterium carotovorum* subsp. *brasiliense* is an emerging SRE with global distribution (35-40). Previous studies have shown that *Pcb* strain PBR1692 (referred to hence forth as *Pcb*1692) is able to inhibit growth of *P. carotovorum* subsp. *carotovorum* WPP14 (*Pcc*) and *P. atrosepticum* SCRI1043 suggesting that many more bacteria can be inhibited by *Pcb*1692 thus giving it a fitness advantage, in a given ecological niche(41, 42). In this study, we use a combination of metagenomics, *in silico* analysis and various *in vitro* and *in planta* competition assays to identify and characterise bacterial targets, and distinctive roles played by the different antimicrobial produced by *Pcb*1692, their regulation and possible contribution towards adaptation to diverse ecological niches.

## Results

### The microbiome of potato tubers

A total of 327,032 and 475,616 paired-end reads from diseased and healthy potato, respectively, were processed. After the initial systematic quality trimming, the 63,841 and 68,015 remaining reads from diseased and healthy samples, respectively, were filtered for chloroplast sequences (see “Methods” for details). A total of 12,379 and 35,526 chloroplast sequences were removed from diseased and healthy samples respectively, and the remaining high-quality reads were analysed using the SILVAngs pipeline (43). Subsequently, a total of 35,177 and 805 reads respectively from diseased and healthy samples were successfully classified into bacterial taxa. The relative distribution of bacteria in healthy and diseased potato tubers is depicted in Fig 1. Overall the analyses showed that representative bacteria from the phylum Proteobacteria were dominant in both diseased (49%) and healthy potato tubers (75%). On the other hand, *Firmicutes* made up 2% and 28% of bacteria in healthy and diseased potato tubers, respectively. Similarly, Gammaproteobacteria were represented in diseased samples mostly by the orders *Pseudomonadales* (25%) and *Enterobacteriales* (56%), in contrast to healthy potato, in which 59% of Gammaproteobacteria were from the *Pseudomonadales* order, and only 31% from *Enterobacteriales* (Fig 1).

**Figure 1.**
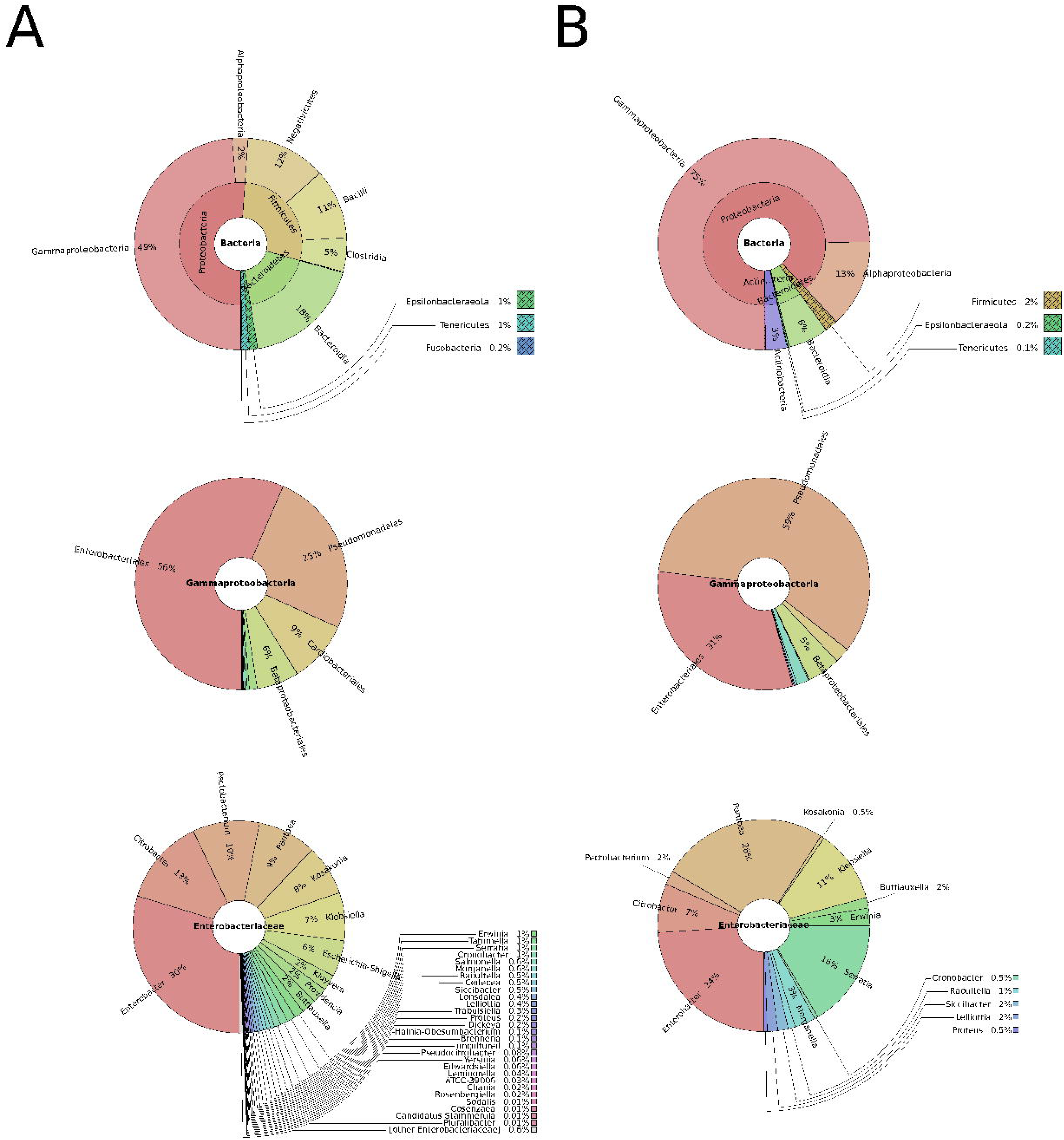
Taxonomic profile of diseased potatoes assessed through shotgun sequencing: Metagenomics analyses represented by composite pie-charts depict the taxonomic representation obtained from sequenced dataset. For each graph, outer radiuses represent the relative proportion of reads assigned to different taxa of the inner circle taxon. Charts under column A represents the relative proportion of reads found in diseased potato tubers while B represents reads from healthy potato tubers.

At a genus level, the analyses identified *Pantoea, Enterobacter, Serratia* together with genera containing important human pathogens such as *Yersinia, Klebsiella, Salmonella, Escherichia, Citrobacter* and *Shigella* in both healthy and diseased potato (Fig 1). Furthermore, soft rot causative pathogens such as *Pectobacterium* and *Dickeya* genera were identified in diseased potato (Fig 1). Importantly, in diseased potato the *Pectobacterium* genus showed a striking 5-fold increase in relative abundance (from 2 to 10% of all *Enterobacteriaceae*) compared to healthy potato tubers (Fig 1).

### Bacterial growth inhibition by *Pcb*1692 is associated with inter and intra species competition

*Pcb*1692 has previously been shown to inhibit growth of some closely related SREs such as *Pcc* and *D. dadantii* (41, 42). In this study we wanted to determine the spectrum of inhibition across different bacteria genera. We hypothesised that *Pcb* strains will likely compete with bacteria that they typically share the same niche. In this regard, bacteria (available from our culture collections) representing selected genera (informed by our metagenomics data) found in healthy and/or diseased potato tubers were selected for subsequent competition assays. Target cells were co-cultured in water or with *Pcb*1692. As has been shown by others, the results show that *Pcb*1692 can inhibit growth of closely related *Pectobacterium* and *Dickeya* species when co-cultured in LB agar (Fig 2). The magnitude of this inhibition is strain specific, as up-to a 6-fold reduction in CFU/ml was observed when *Pcb*1692 was co-cultured with some members of SRE such as *P. atrosepticum, Pcc, Pectobacterium carotovorum subsp.odoriferum* (*Pco*) and *D. dadantii* while moderate levels of inhibition 1-3 fold reduction was observed for *P. paradisiaca, P. cipripedii, P. cacticida and P. betavasculorum.* The study also demonstrates that *Pcb*1692 is also able to outcompete bacteria other than those within the SRE such as *Enterobacter cowanii, Pseudomonas aeruginosa, Salmonella typhimurium, Escherichia coli* and *Serratia marcescens* (Fig 2). The ability of *Pcb* to inhibit *P. aeruginosa, S. typhimurium, E. coli, Enterobacter cowanii* and *S. marcescens*, which we identified as endophytes of potato tubers likely gives *Pcb* a fitness advantage in potato tubers. Interestingly, the results also showed that *Pcb*1692 was able to inhibit growth of other *Pcb* strains G4P5, G4P7, XT3 and XT10 but not strains HPI01, 358, CCI and CC2 (Fig 2, and results not shown). The reason for this intraspecies inhibition is unclear but it is possible that *Pcb*1692 secretes some yet to be identified effectors for which some closely related strains lack the immunity factor needed to neutralize its cytotoxic effect. In addition, the data also showed that *Pcb*1692 was unable to inhibit *in vitro* growth of bacteria such as *D. chrysanthemi, P. wasabiae, S. fiscaria, P. ananatis, P. stewartii* subsp. *indologenes, Bacillus subtilis* and *B. cereus* in rich LB medium. Together, these findings demonstrate that, while *Pcb*1692 readily inhibits growth of most SREs analyzed in this study, including closely related *Pcb* strains, it also inhibits growth of non-SREs.

**Figure 2.**
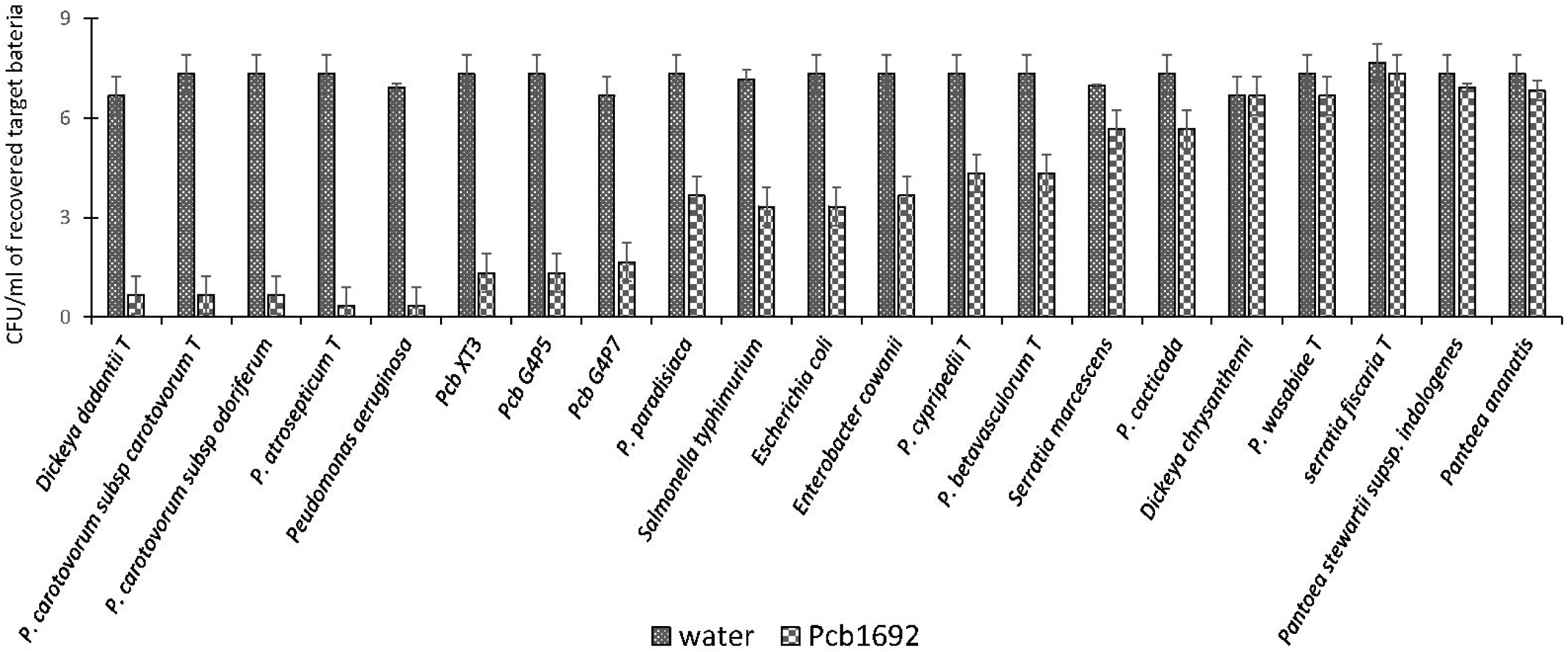
Interbacterial competition between *Pectobacterium carotovorum* subsp. *brasiliense* (*Pcb*1692) and targeted bacteria: *Pcb*1692 and targeted bacteria were cocultured in a 1:1 ratio on LB and the CFU/ml of surviving targeted bacteria enumerated 24hrs post incubation. Experiments were performed in triplicates and repeated three independent times.

### Iron in potato tuber extracts plays an essential role in bacterial competition

Potato tubers contain several metals such as iron, potassium, zinc, copper, manganese, lead, cadmium with the highest concentrations being potassium and iron (44, 45). Some of these metals play a role in virulence and bacterial competition, by either acting as enzyme co-factors or co-activator/-repressors of some transcriptional regulators (46). We therefore investigated the effect of some of these metals on the observed ability of *Pcb*1692 to kill targeted bacteria *in vitro*. For this part of the study, *Pcb*1692 was co-cultured with *Pcc* and *D. dadantii* in M9 minimal media supplemented with different metals (see Materials and Methods). The results showed that while low concentrations of copper and cobalt (0.01 µM and 1µM) had no effect on competition, higher concentrations of both metals (10µM and 50 µM) inhibited growth of both *Pcb*1692 and target bacteria (results not shown). In addition, different concentrations of magnesium, manganese, zinc, nickel (0.01, 1, 10 and 50 µM) had no effect on bacterial competition (results not shown). On the other hand, the addition of ferric iron (10 or 50µM) to M9 minimal medium enhanced the ability of *Pcb*1692 to kill either *Pcc, Pco* or *D. dadantii* while lower concentrations (0.1 and 1 µM) had no effect on bacteria competition (Fig 3 and results not shown). Therefore, 10 µM ferric iron was used to supplement M9 medium in all downstream assays associated with iron. For competition assays, target bacteria were co-cultured in water (controls) or with *Pcb*1692 (Fig 3). Figure 3 shows that when *Pcc* or *Pco* were co-cultured with *Pcb*1692 in LB (rich) medium, fewer cells of target bacteria were recovered compared to controls. However, a 1-2 fold increase in the number of target bacteria cells recovered after co-culture in nutrient-poor M9 medium (relative to rich medium) was observed. This indicated reduced *Pcb*1692-killing ability in nutrient-poor medium compared to LB rich medium. The magnitude of inhibition by *Pcb*1692 increased 1-2 folds when M9 agar was supplemented with potato tuber extracts. Thus, adding tuber extracts to minimal medium enriched the medium and hence a similar outcome on competition as with rich LB medium was observed. When M9 was supplemented with iron, fewer competitor/target cells were recovered (relative to rich medium), indicating increased *Pcb*1692-killing ability. Addition of the iron chelator 2,2’-dipyridy to M9 tuber-supplemented medium eliminated ability of *Pcb*1692 to kill competing bacteria, while addition of 10µM of ferric iron to M9 media restored competition in the absence of plant extracts. Together these findings demonstrate that the presence of iron increased the ability of *Pcb*1692 to inhibit targeted bacteria. In the next sections we report the approaches used to establish whether or not the effect of iron is associated with the expression of T6SS, carbapenem or bacteriocin production in *Pcb*1692.

**Figure 3.**
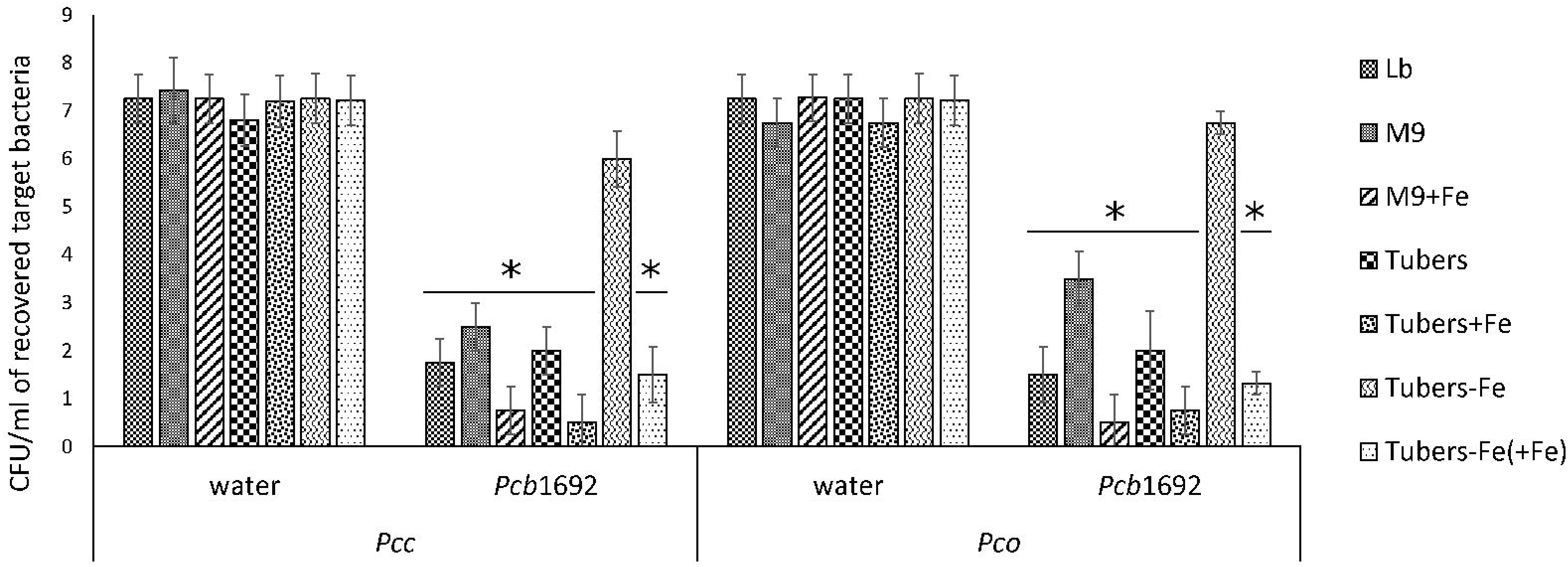
Effect of iron in interbacterial competition: *Pectobacterium carotovorum* subsp. *brasiliense* 1692 and targeted bacteria were cocultured in a 1:1 ratio in either M9, M9 supplemented with either potato tuber extracts or 10µM ferric iron. The iron in plant extracts was removed using 2,2’dipyridy. Results are presented as the CFU/ml of recovered targeted bacteria following 24hr co-culture. Asterisk indicates statistically significant differences *p*<0.05 relative to water control and targeted strains co-cultured with *Pcb*1692 on agar containing potato extracts pre-treated with 2,2’dipyridy. *Pcc* = *Pectobacterium carotovorum* subsp. *carotovorum, Pco* = *Pectobacterium carotovorum* subsp. *odoriferum*.

### *In silico* identification of genes associated with antibacterial activity

A transcriptome profiling experiment performed previously in our laboratory, showed that *Pcb*1692 genes encoding the T6SS and bacteriocins (carotovoricin and pyocin) were up-regulated during *in planta* (potato tuber) infection (47). The same study also showed that gene homologs of the *Pcb*1692 T6SS were found to be present in the genome sequences of all publicly available genome sequences of *Dickeya* and *Pectobacterium* (47). Similarly, while homologous sequences of *Pcb*1692 carotovoricin gene cluster were present in all publicly available genome sequences of *Pectobacterium* spp. it is missing from the genome sequences of all *Dickeya* spp. (except *D. dianthicola* RNS049 and *D. zeae* strains CSLRW192, EC1 and ZJU1202). The above analysis was extended to include orthology relationship between the *Pcb*1692 protein sequences associated with the carbapenem and the S-type pyocin in order to determine their presence or absence in other *Pectobacterium* and *Dickeya* spp.

The *Pcb*1692 *car* gene cluster was found to contain all six conserved genes required for synthesis of carbapenem (*car*A/B/C/D/E/H) including genes encoding the carbapenem immunity proteins (CarF/G). Homologous *car* gene clusters were also identified in *P. betavasculorum* ATCC 43762 (T), *Pcc* ATCC 15713(T), *D. chrysanthemi* NCPPB 3533, *D. zeae* CSL RW192, *D. zeae* Ech1591 and several strains of *Pcb* (Fig 4). This gene cluster was found to be missing from all publicly available genome sequences of *P. wasabiae, P. atrosepticum, P. parmentieri, D. solani* and *D. dadantii* (exempting *D. dadantii* subsp. *dieffenbachiae* NCPPB2976) (S1 Table-Sheet1). Similarly, the *car* gene cluster was found to be missing from the genome sequence of *Pcb* strains (BC1, YCD21, YCD49, YCD62, and YCD65). Interestingly, although lacking the *car* gene cluster, *P. wasabiae* strains (CFBP3304, NCPPB3701, and NCPPB3702), *P. parmentieri* strains (CFIA1002 and RNS08421a) and *Pcb* strains (BC1, YCD21, YCD49, YCD62, and YCD65) all retained the carbapenem immunity gene *car*F*/car*G, suggesting that they may be resistant to the carbapenem produced by some *Pectobacterium* and *Dickeya* spp.

**Figure 4.**
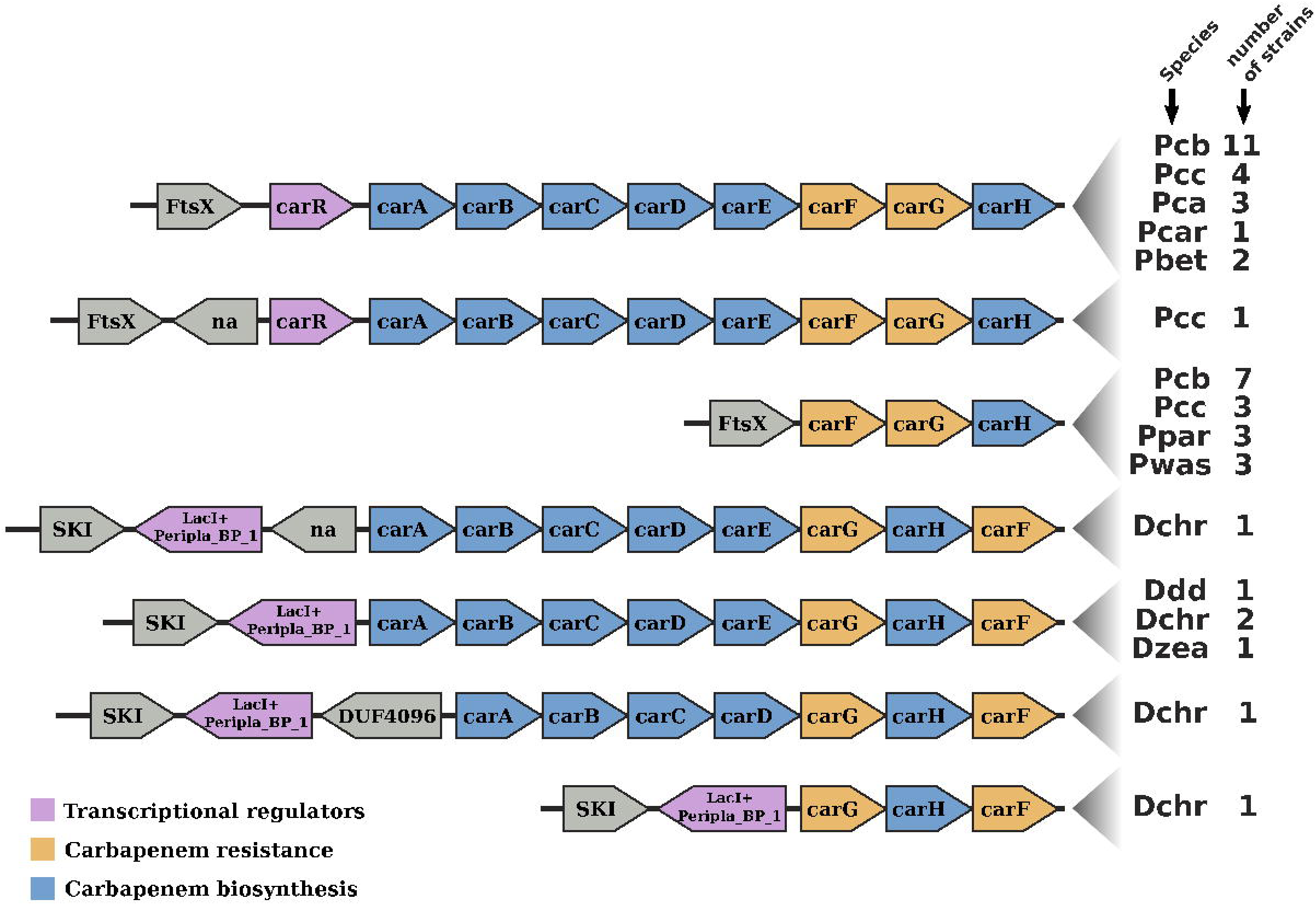
Gene-neighborhood screening of the carbapenem gene cluster (*car*) in Soft-Rott-Enterobacteriacea (SRE): 100 SRE genomes were analysed and all bacteria strains that conserve segments of the *car* cluster were screened. All known *car* genes are labelled in the figure. Genes unrelated with the *car* cluster are labelled using functional domains detected in the respective encoded proteins when applicable, otherwise these are labelled with “na”.

Furthermore, we identified a putative S-type pyocin encoded by the *Pcb*1692 *PCBA_RS02805*. This protein (794 amino acids) was predicted to contain a colicin-like bacteriocin tRNase domain (Pfam: PF03515)(48), suggesting it might kill other bacteria by degrading their tRNA. Homologs of the *Pcb*1692 pyocin were identified in the genome sequence of *Pcb* strains (LMG21371, HPI01, YCD29, ICMP19477, BC1), *Pcc* strains (PCC21, NCPPB3395 and YCD57) and *P. betavasculorum* NCPPB2795 but missing from all publicly available genome sequences of *P. atrosepticum, P. wasabiae* and *Dickeya* (S1 Table-Sheet-2). Downstream of the *Pcb*1692 *PCBA_RS02805* gene is *PCBA_RS02810* encoding the cognate S-pyocin immunity factor (∼90 amino acids), with homologs present in all SREs encoding S-pyocin. Interestingly, although the genome sequences of *Pc. odoriferum* (*Pco*) strains (S6, S6-2, Q106, Q32, and Q3) and *Pcc* strains (Y57 and 3F-3) do not have genes encoding the homologous *Pcb*1692 S-pyocin, they encode S-pyocin immunity factors 100% identical to the one found in *Pcb*1692 (PCBA_RS02810). Overall, this analysis demonstrates that the *Pcb*1692 toxin/antitoxin modules analysed in this study are not conserved in all bacteria including some strains of *Pcb* and may therefore be a determining factor in the outcome of inter or intra species competition among SRE.

### The *Pcb*1692 carbapenem gene cluster plays a role in bacterial competition *in vitro*

In order to investigate the relative contribution of the *Pcb*1692 T6SS, carotovoricin, carbapenem and the putative pyocin in competition, we generated *Pcb*1692 mutants with deletions in genes associated with the production of S-pyocin (*Pcb*1692Δ*pyo*), carbapenem (*Pcb*1692Δ*carC*), carotovoricin (*Pcb*1692Δ*lyt* and *Pcb*1692Δ*fer*) and biosynthesis of a functional T6SS (*Pcb*1692ΔT6). Targeted bacteria (*D. dadantii* or *Pcc*), earlier shown to be inhibited by wild-type *Pcb*1692, were co-cultured *in vitro* with either water or the different *Pcb*1692 mutant strains. The results show that the CFU/ml of recovered targeted bacteria following co-culture with the *Pcb*ΔT6, Δ*lyt*, Δ*fer* and Δ*pyo* mutant strains was similar to the CFU/ml of recovered targeted bacteria following co-culture with the wild-type strain. However, the *Pcb*1692Δ*car*C mutant strain lost the ability to inhibit growth of *D. dadantii* and *Pcc* (Fig 5A). Trans-expression of the *car*C gene in *Pcb*1692Δ*car*C restored the ability of the complemented strain to inhibit targeted bacteria, similar to wild-type *Pcb*1692. (Fig 5A). Together, these findings suggest that carbapenem production by *Pcb*1692 is associated with the interbacterial and intrabacterial competition *in vitro*. Trans-expression of the *Pcb*1692 *car*F/G genes in *D. dadantii* prevented growth inhibition of this strain when co-cultured with wild-type *Pcb*1692 confirming that growth inhibition of these bacteria is associated with *Pcb*1692 carbapenem and that the *Pcb*1692 *car*F/G genes confer immunity to the *Pcb*1692 carbapenem (Fig 5Bi and ii). In addition, our results showed that while *Pcb* strains CC1, CC2, HPI01 and 358 can produce carbapenem, which inhibit growth of *D. dadantii, Pcb* strains XT3, XT10, G4P5 and G4P7 do not produce carbapenem under the same experimental conditions (Fig 6). Similarly, the carbapenem produced by *Pcb* strains 1692, CC1, CC2, HPI01 and 358 was found to inhibit growth of *Pcb* strains XT3, XT10, G4P5 and G4P7, suggesting that the latter group of strains lack the cognate carbapenem immunity factor. Together, *in vitro* and *in silico* analyses uncover the contrasting conservation patterns of the carbapenem biosynthesis cluster in SREs, highlighting its critical role in interference competition among these organisms. With this information, we proceed toward assessing possible regulatory processes involved in the control of carbapenem biosynthesis.

**Figure 5.**
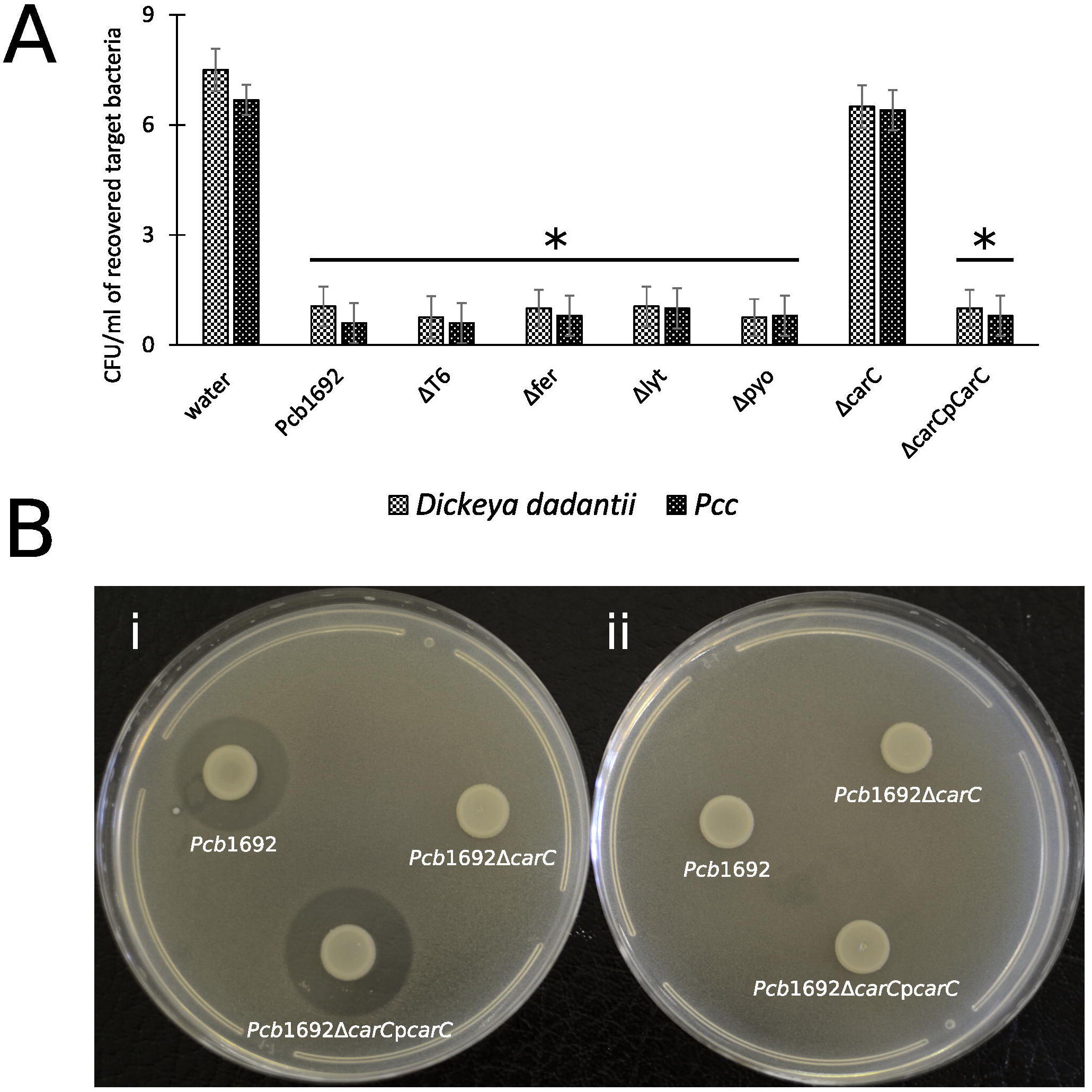
Interbacterial competition with *Pectobacterium carotovorum* subsp. *brasiliense* 1692 mutant strains. **A)** Wild type *Pcb*1692 and its isogenic mutant (T6, *fer, lyt*, and *carC*) were cocultured with *D. dadantii* and *Pcc* for 24hrs. The data shows the CFU/ml of recovered *D. dadantii* and *Pcc* following co-culture. The horizontal line with an asterisk represents the range of samples with statistically significant differences (*p*<0.05) relative to water controls. B) Bi) Clearing zone die to carbapenem produced by strains of *Pcb*1692 spotted on a lawn of *D. dadantii* with circularize plasmid PJET4, Bii) No clearing zones when of *Pcb*1692 is spotted on a lawn of *D. dadantii* expressing *car*F and G resistance genes from plasmid pJET4-*car*FG.

**Figure 6.**
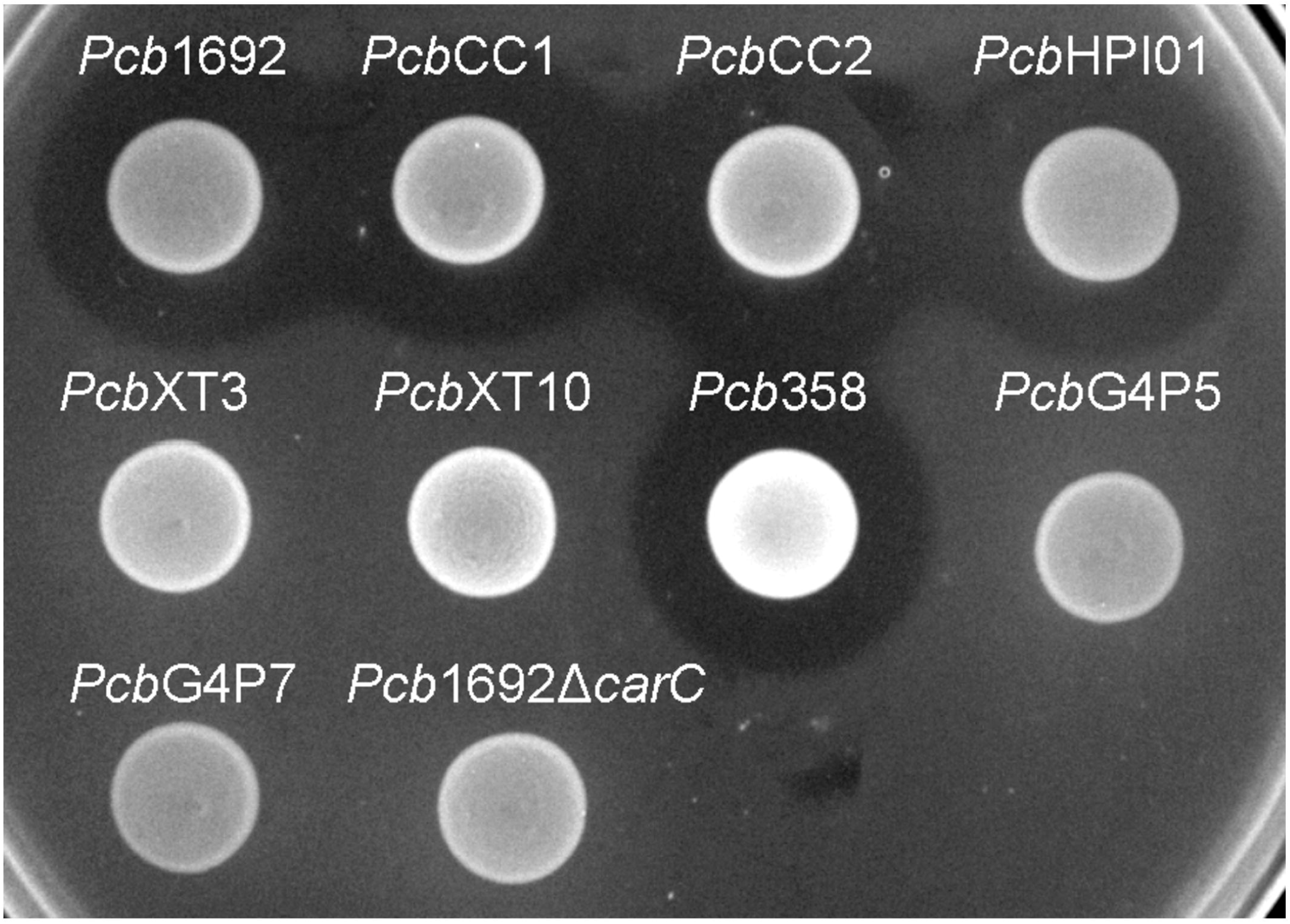
Carbapenem production by environmental strains of *Pectobacterium carotovorum* subsp. *brasiliense.* The ability of laboratory strains of *Pcb* isolated from South Africa to produce carbapenem was determined by the spot-on-lawn assay, as previously described. The presence of a clear zone around the spotted *Pcb* was indicative of carbapenem production. *Pcb*1692 was used as a control given that this strain produces carbapenem.

### Iron is required for carbapenem production by *Pcb*1692 in minimal media and the *car* gene cluster is regulated by Fur, SlyA and ExpI

Having demonstrated that the *Pcb*1692 carbapenem plays a role in *in vitro* interbacterial competition, we next investigated how this gene cluster is regulated and under which conditions *Pcb*1692 produces carbapenem. To this end, we included *Pcb*1692 transcriptional regulator mutant strains generated from previous studies in our laboratory including the ferric uptake regulator (Fur), N-acyl homoserine lactones (18) synthase (ExpI) and the stress response regulator SlyA. The results showed that while wild-type *Pcb*1692 produced a clear zone of inhibition when cultured on a lawn of *P. atrosepticum*, the *Pcb* mutant strains (Δ*car*C, Δ*sly*A, Δ*exp*I and Δ*fur*) did not produce this clear zone associated with carbapenem production (Fig 7A). Loss of carbapenem production could be complemented by trans-expression of corresponding genes in the *Pcb*1692 mutant strains (Fig 7A). In addition, while wild-type *Pcb*1692 and the complemented mutant strains produced carbapenem on LB, they all lost the ability to produce carbapenem on M9 supplemented with 0.4% of glucose (Fig 7B). Interestingly, carbapenem production was restored when M9 was supplemented with 10 µM of ferric iron [results not shown]. These findings implicate iron and the iron homoeostasis protein Fur in the regulation of carbapenem production in *Pcb*1692. Aiming to predict whether Fur is able to regulate carbapenem production in *Pcb*1692 directly, or indirectly, we performed *in silico* analysis using the MEME package (49) to search for *fur* binding sites in the promoter regions of *car*R, *sly*A, *exp*I and *exp*R. Interestingly, the analysis identified *fur* binding sites in the promoter sequences of *car*R, *sly*A and *exp*R but not in the promoter sequences of *exp*I and *car*A (the first gene in the *car* gene cluster) (S2 Table). Given that CarR, and not ExpR, is required for carbapenem production (26, 50, 51), the *Pcb*1692 Fur protein may be able to indirectly control the transcription of carbapenem production via *car*R, *sly*A, or both.

**Figure 7.**
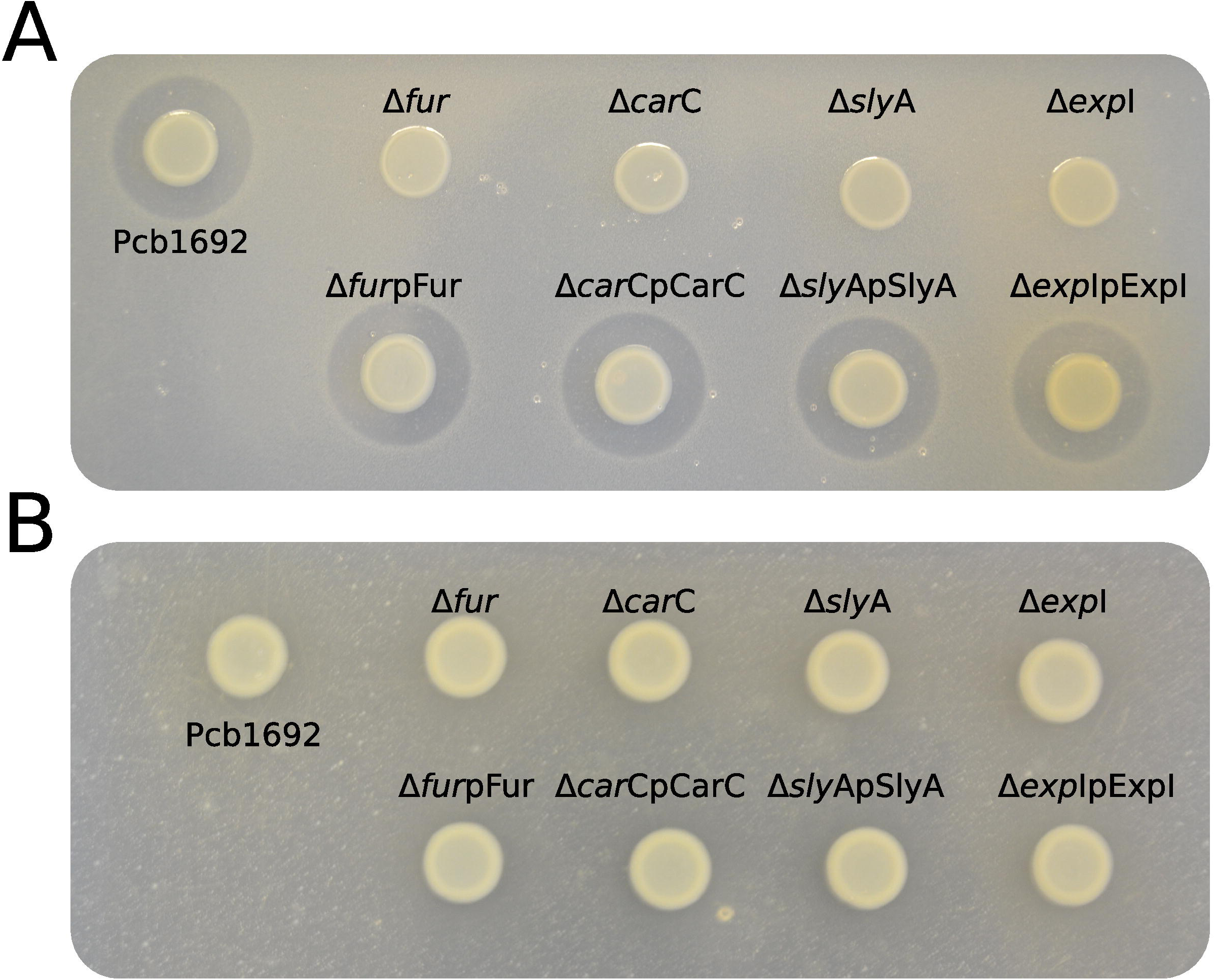
Iron is required for production of carbapenem by *Pectobacterium carotovorum* subsp. *brasiliense* 1692. *Pcb1692* wild type mutant and complemented mutant strains were spotted on a lawn of *P. atrosepticum* and clear zone around the spotted *Pcb* strains was indicative of carbapenem production. A) carbapenem production on LB agar by *Pcb* strains B) No detectable levels of carbapenem were observed on M9.

### The *Pcb*1692 T6SS plays a role in interspecies but not intraspecies competition *in planta*

Given the *Pcb*1692 T6SS did not appear to play a role in bacterial competition *in vitro*, but was upregulated in potato tubers, we reasoned that *Pcb*1692 T6SS could play a role in competition *in planta*. To determine the contribution of the *Pcb*1692 T6SS in bacterial competition, *in planta*, standardized cultures of *Pcb*1692, *Pcb*1692ΔT6 or *Pcb*1692ΔT6-pT6 were independently co-inoculated with targeted bacteria (*D. chrysanthemi, D. dadantii, Pcc, P. atrosepticum* or *Pcb*G4P5) into surface sterilized potato tubers, and CFU/ml of targeted bacteria enumerated three days post-inoculation. The results show no reduction in the CFU/ml of *Pcb*G4P5 when co-inoculated in potato tubers with either wild-type *Pcb*1692, *Pcb*1692ΔT6 or *Pcb*1692ΔT6-pT6 suggesting that effectors secreted by the T6SS of *Pcb*1692 cannot inhibit growth of this particular strain [results not shown]. However, a threefold reduction in the CFU/ml was observed when *D. dadantii, Pcc, P. atrosepticum* or *D. chrysanthemi* were co-inoculated with the wild-type *Pcb*1692 and complement, *Pcb*1692ΔT6-pT6, this inhibition was completely lost when targeted strains were co-inoculated with *Pcb*1692ΔT6 (Fig 8 and results not shown). These results demonstrate the involvement of the *Pcb*1692 T6SS in interbacterial competition in potato tubers. We earlier demonstrated that although the *in vitro* production of carbapenem by *Pcb*1692 plays a role in both intra and interspecies competition, it does not inhibit growth of *D. chrysanthemi*. The *D. chrysanthemi* strain used in this study was found to be resistant to ampicillin (results not shown), suggesting that this strain might produce a β-lactamase which degrades carbapenem. Conversely, although the results have not been shown, we found that the T6SS of *Pcb*1692 inhibits growth of *D. chrysanthemi in planta* but not *in vitro*. These findings together with our *in vitro* competition assays suggest that depending on the ecological niche, *Pcb*1692 can use either the T6SS or carbapenem to inhibit growth of several bacteria species. This is further supported by the fact that the *Pcb*1692 *car* gene cluster, unlike the T6SS, is not differentially expressed *in planta* and may thus be recruited by *Pcb*1692 to eliminate competitors while residing on the rhizosphere or plant surfaces (47). However, following ingress into plant tissue *Pcb*1692 likely recruits mainly the T6SS for bacterial competition.

**Figure 8.**
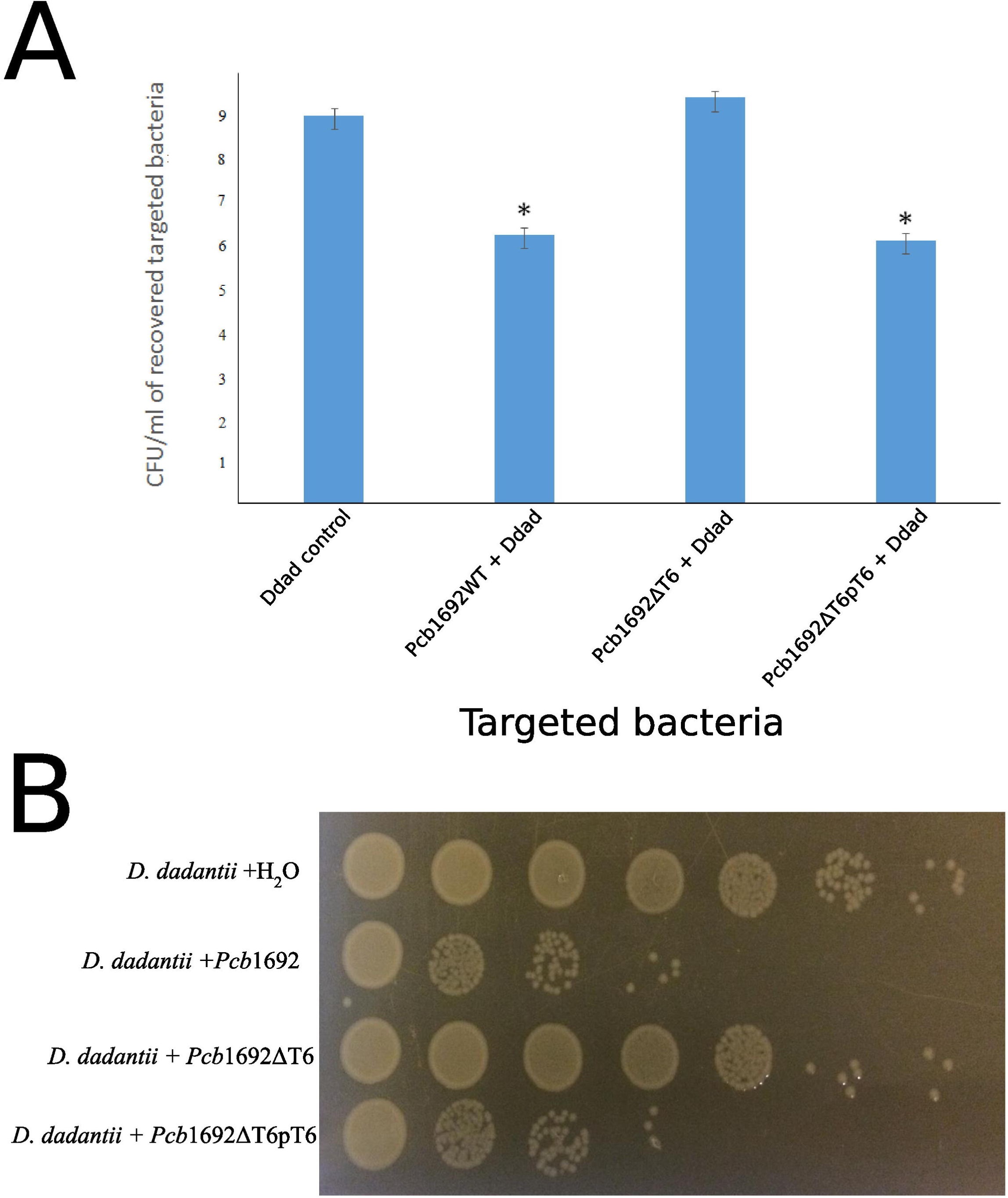
Effect of the T6SS in *Pcb*1692 anti-bacterial competition *in planta*. Susceptible potato tubers (*S. tuberosum* cv. Mondial) were co-inoculated with standardized cultures mixed in a 1:1 ratio of *Dickeya dadantii* (gentamycin resistant) with either water, wildtype *Pcb*1692, *Pcb*1692ΔT6 *or Pcb1692*ΔT6pT6, incubated for 72hr and A) survival of *D. dadantii* enumerated by serial dilution and recorded as CFU/ml. D. dad = *D. dadantii* b) recovered of *D. dadantii* on LB supplemented with 15µg/ml of gentamycin. Asterisks indicated statistically significant differences (*p*<0.05) relative to water controls.

### *Pcb*1692 produces bacteriocin(s) following mitomycin C induction

Despite the fact that the genome sequence of *Pcb*1692 has genes encoding pyocin and carotovoricin, no role was determined for these bacteriocins in the assays we conducted thus far. Bacteriocin production is usually induced in response to DNA damaging agents which trigger SOS response (52, 53). Thus we investigated the ability of *Pcb*1692 to produce these bacteriocins following induction with either mitomycin C or UV irradiation, using the spot-on-lawn overlay method (see Materials and Methods for details). The results showed that wild-type *Pcb*1692, *Pcb*1692Δ*car, Pcb*1692Δ*fur, Pcb*1692Δ*sly*A and *Pcb*1692Δ*exp*I all produced bacteriocins following mitomycin C treatment and no bacteriocins were produced in control experiments (Fig 9A and B and results not shown). Considering that the above mentioned *Pcb*1692 mutant strains do not produce carbapenem, this indicated that the clear zones around the spotted bacteria may either represent *Pcb*1692 S-type pyocin, carotovoricin, a combination of both bacteriocins or a yet to be identified bacteriocin. In order to determine whether these clear zones were due to secretion of the putative S-type pyocin by *Pcb*1692, we generated a *Pcb*1692Δ*pyo*I double mutant strain lacking both the pyocin and putative immunity genes but with intact genes associated with carbapenem and carotovoricin production. Interestingly, when *Pcb*1692Δ*pyo*I was overlaid on mitomycin C-induced *Pcb*1692 strains, we observed clear zones of inhibition around wild-type *Pcb*1692 and all mutant strains used in study except *Pcb*1692Δ*pyo*, which lacks the S-type pyocin but has the immunity gene (Fig 10). In addition, the complemented pyocin mutant strain *Pcb*1692Δ*pyo*p*pyo* was restored in its ability to inhibit growth of *Pcb*1692Δ*pyo*I similar to wild-type *Pcb*1692 while no *Pcb*1692 strain could kill the complemented *pyo*I mutant *Pcb*1692Δ*pyo*Ip*pyo*I (Fig 10). Together these findings demonstrate that following mitomycin C induction, *Pcb*1692 secretes the S-type pyocin, which inhibits growth of bacteria lacking the immunity factor. To evaluate, host range of killing by the *Pcb*1692 S-type pyocin, the overlay assay was repeated with different bacteria and the results show that bacteriocins produced by *Pcb*1692 can inhibit growth of *P. atrosepticum, Pcc, Pco, D. dadantii, S. typhimurium* and *E. coli* but not the type strains of *S. marcescens* and *D. chrysanthemi* including some environmental strains of *Pcc* isolated from South Africa (S3 Table). The results also showed that the *Pcb*1692 carotovoricin isogenic mutant strains *Pcb*1692Δ*fer* and *Pcb*1692Δ*lyt* produced bacteriocins similar to the wild-type. This could be because the *Pcb*1692 carotovoricin mutant strains still produced S-type pyocin or because the ferredoxin gene located within the *ctv* gene cluster of *Pcb*1692 may not be essential for carotovoricin production given that it is not conserved in genome sequence of several SREs that have this gene cluster (47). Similarly, the carotovoricin lytic cassette has three lytic genes associated with cell lysis and release of carotovoricin, thus presenting the possibility of functional complementation following deletion of any one the lytic genes, in our case, *Pcb*1692 Δ*lyt*.

**Figure 9.**
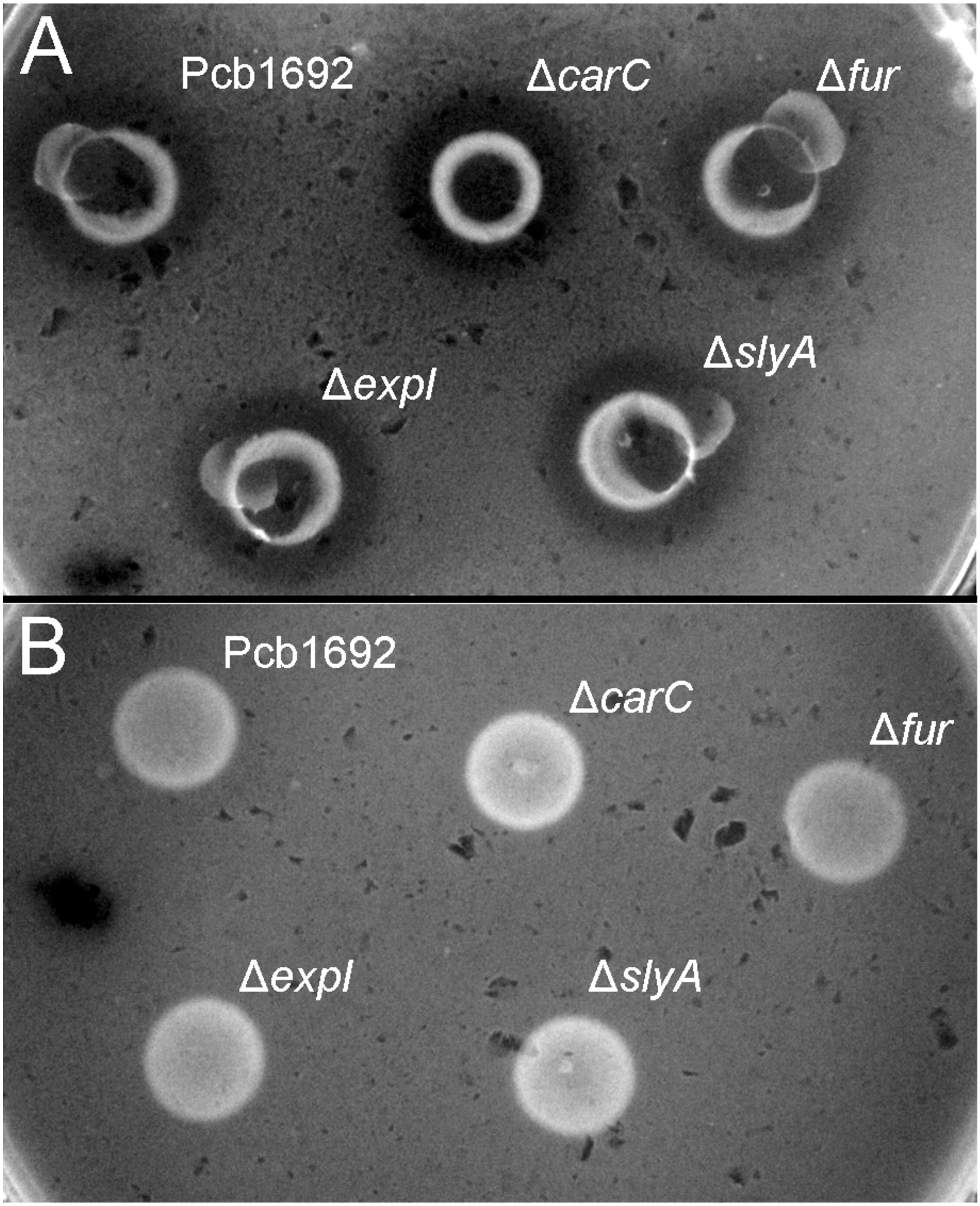
Bacteriocin production by strains of *Pectobacterium carotovorum* subsp. *brasiliense* 1692. *Pcb*1692 and its isogenic mutant strains were spotted on LB and grown overnight. A) bacteriocin production was induced with mitomycin C B) control plate not induced. All plates induced and untreated control were incubated for 5hrs, kill with chloroform vapour and overlaid with a lawn *Dickeya dadantii*. Clear zones are indicative of bacteriocin production.

**Figure 10.**
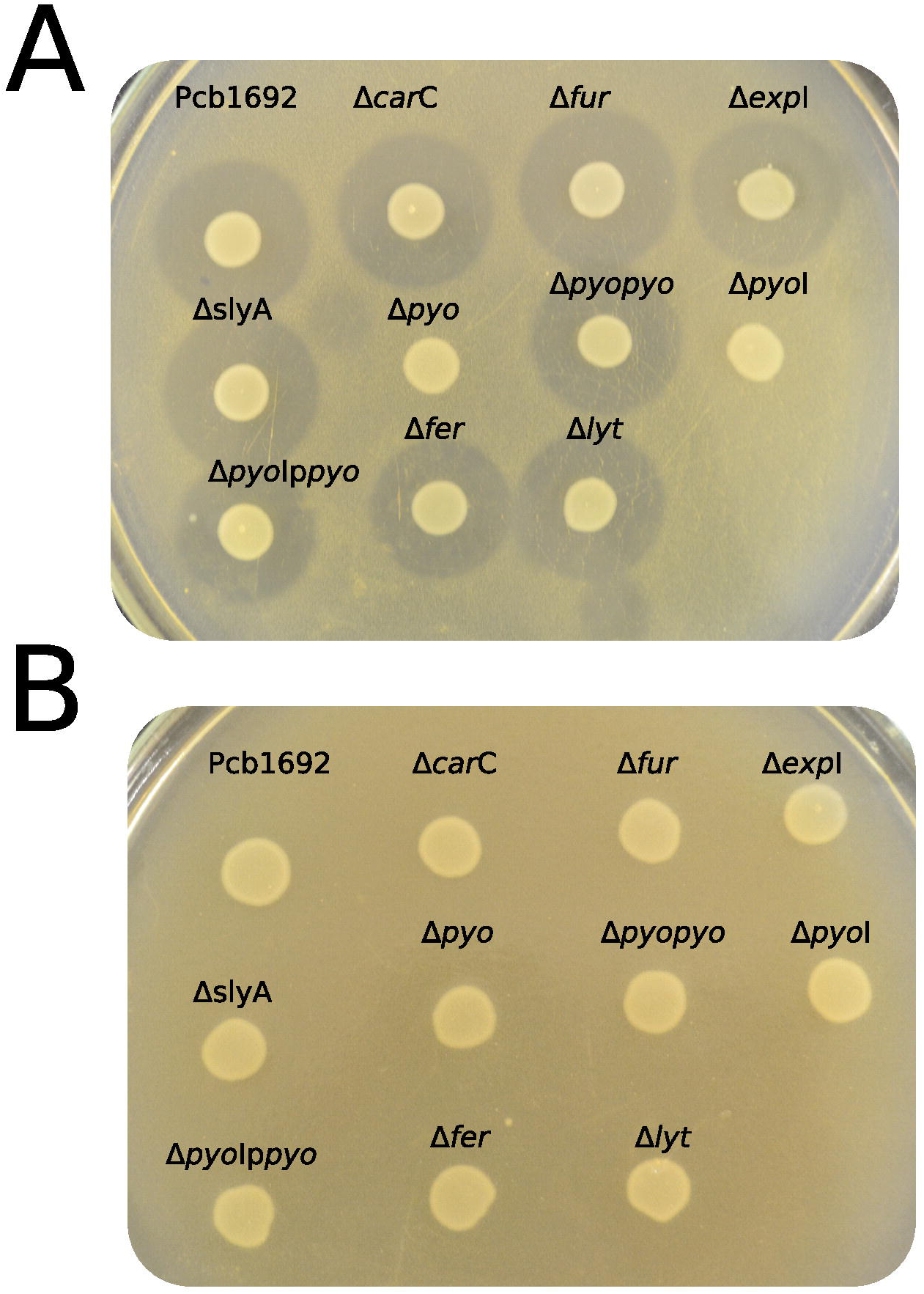
Production and detection of the S-type pyosin of *Pectobacterium carotovorum* subsp. *brasiliense* 1692. *Pcb*1692 and its isogenic mutant strains including *Pcb*1692Δpyo, *Pcb*1692ΔpyoI and their respective complemented strains, were spotted on LB and grown overnight. Bacteriocin production was induced with mitomycin C for 5hrs. Induced cells were killed with chloroform vapour and overlaid with A) a lawn of either *Pcb*1692Δ*pyoI* or *Pcb*1692Δ*pyoI*p*pyoI*. Clear zones are indicative of bacteriocin production.

### *Pcb*1692 produces bacteriocins but not carbapenem under anaerobic conditions

Previous studies have shown that oxygen is required for carbapenem production (26, 54). Therefore, to better understand the role played by oxygen in bacterial competition and the biological significance of carbapenem and bacteriocin production in the context of potato tubers, we assayed the ability of *Pcb*1692 to produce these antimicrobial compounds under aerobic or anaerobic conditions. The results demonstrated that *Pcb*1692 produces carbapenem under aerobic but not anaerobic conditions (Fig 11A). In addition, following mitomycin C treatment *Pcb*1692 and its isogenic mutant strains *Pcb*1692Δ*car*C, *Pcb*1692ΔT6, *Pcb*1692Δ*pyo, Pcb*1692Δ*fer* and *Pcb*1692Δ*lyt* all produced bacteriocins when overlaid with either *Pcc, Pco, P. atrosepticum, D. dadantii* or *Bacillus cereus* under both aerobic and anaerobic conditions (Fig11B, Supple Table2 and results not shown). We further investigated the role played by the *Pcb*1692 T6SS in competition under anaerobic conditions, which mimics one of the conditions *in planta* where there is limited oxygen. Under aerobic conditions, no targeted bacteria were recovered when co-cultured with either *Pcb*1692 wild type, mutant or complemented mutant strains. This could be ascribed to previously demonstrated production of *Pcb*1692 carbapenem under aerobic conditions (Fig 11A). *Pcb*1692 killing ability on target bacteria under anaerobic conditions was only slightly reversed in the *Pcb*1692ΔT6 mutant strain. Thus, we could conclude that the presence or absence of oxygen did not affect T6SS dependent *in vitro* bacterial competition (Fig11C). Together these findings suggest that *Pcb*1692 might secrete bacteriocins but not carbapenem following ingress into potato tubers, where conditions are mostly anaerobic. Furthermore, the presence or absence of oxygen did not affect the magnitude of growth inhibition of targeted bacteria by *Pcb*1692ΔT6 relative to wild-type. This therefore supports our previous findings that the T6SS of *Pcb*1692 may only be functionally active *in planta*.

**Figure 11.**
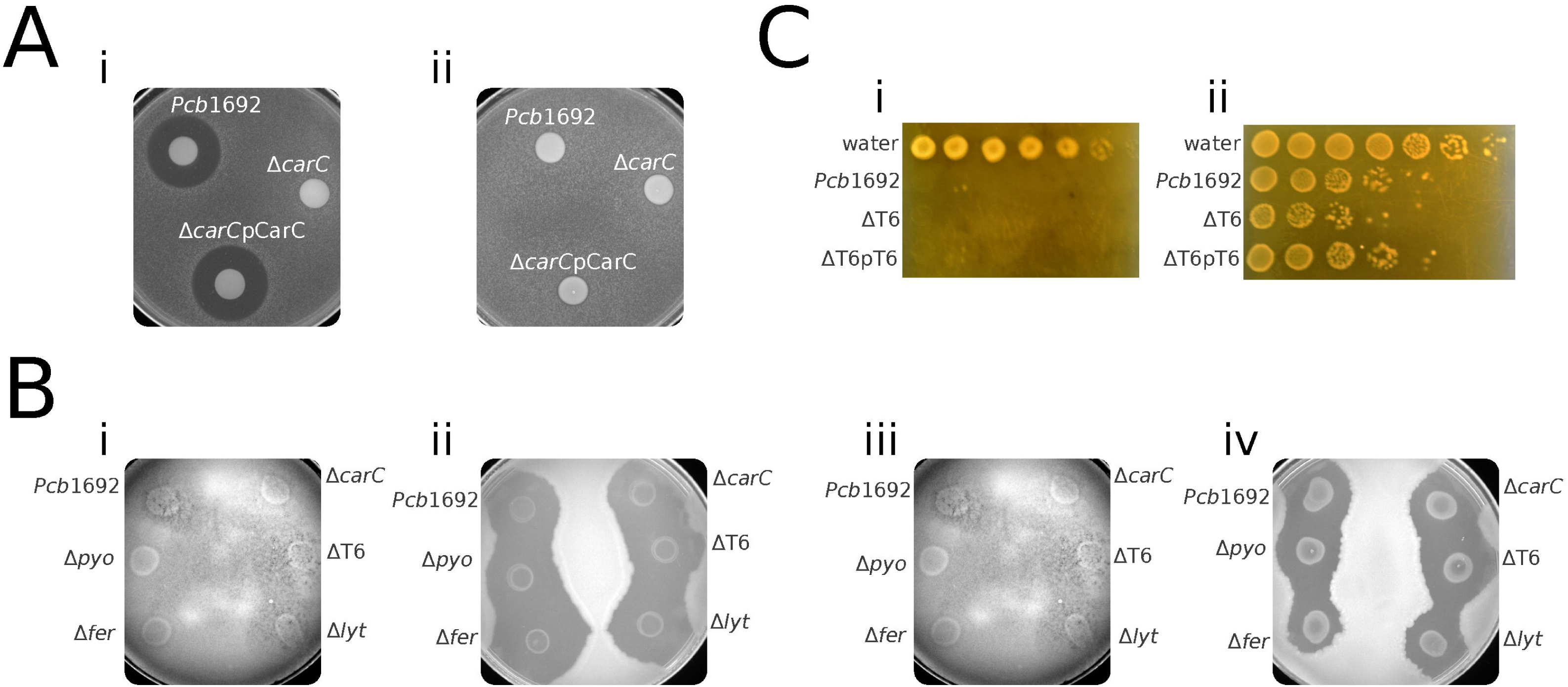
Effect of oxygen in competition, antibiotic and bacteriocin production by *Pectobacterium carotovorum* subsp. *brasiliense* 1692. A) carbapenem production by *Pcb*1692 strains incubated i) aerobically, without prior induction (control) and ii) anaerobically induced with mitomycin C B) Bacteriocin production by *Pcb*1692 strains incubated i) aerobically without prior induction ii) aerobically induced with mitomycin C, iii) anaerobically without induction (control) iv) anaerobically with mitomycin C induction and Interbacterial competition assays between *Pcb*1692 and the targeted strain *Pcb*G4P5, data is presented as CFU/ml of recovered targeted bacteria following 24hr incubation either i) aerobically or ii) anaerobically.

## Discussion

Bacteria exist as complex multispecies communities in any given niche (7, 10). The interactions within these communities can be characterised by cooperation between species or more commonly, by competition (55, 56). We set out to explore microbe-microbe interactions that shape population structures of soft rot *Enterobacteriaceae*. We used *Pcb*1692, an emerging soft rot pathogen of potatoes, as a model to help us understand factors shaping dynamics of microbial communities found in decaying potato tubers.

In the metagenomics screening, population shifts from uninfected to infected potatoes showed that *Pseudomonads* were more represented in healthy potato than in diseased potato while *Firmicutes* and several *Proteobacteria* exhibited stronger representation in diseased potato. It is known that some *Firmicutes* belonging to the species *Bacillus* and *Clostridium* exhibit amylase and pectolytic activity, which gives them the ability to take up nutrients which are not readily available to other endophytes (33, 57, 58). However, once SREs and *Firmicutes* break down plant cell wall, nutrients readily become available to the fast growing *Proteobacteria,* which outcompete *Pseudomonads*. In the early stages of disease development, endophytic SREs and *Firmicutes* may cooperate to degrade plant cell wall to acquire nutrients, however, in the absence of disease as is the case in healthy potato, these latent pathogens will have to compete for space, water and limited nutrients which could result in an antagonistic interaction.

Interference competition is characterised by deployment of antibacterial weapons to eliminate target bacteria (56, 59). The *in vitro* competition assays in LB rich media showed that *Pcb*1692 inhibits growth of several bacteria such as *P. atrosepticum, Pcc, Pco, D. dadantii, Serratia marcescens, Salmonella typhimurium, E. coli* and several laboratory strains of *Pcb*, but not *P. wasabiae, D. chrysanthemi* and *Pantoea ananatis, B. subtilis and B. cereus*. To do this, *Pcb* recruits an impressive number of antibacterial weapons. Among these, is *Pcb*1692 T6SS which inhibits growth *D. dadantii, Pcc* and *D. chrysanthemi* but not environmental strains of *Pcb* when co-inoculated in potato tubers. This T6SS-mediated inhibition was completely lost when competition assays were performed *in vitro*. In contrast, it has been previously reported that the T6SS of *Pcb*1692 is upregulated *in planta* (47), suggesting that this system is only functional following perception of an unknown potato tuber signal which may have been lost during preparation of plant extracts (60-64).

Furthermore, *Pcb*1692 deploys carbapenem and bacteriocins to inhibit growth of all the aforementioned targeted bacteria, except *Serratia marcescens, D. chrysanthemi* and *P. wasabiae*. The results show that under anaerobic conditions *Pcb*1692 produces detectable levels of carbapenem and bacteriocins when spotted on a lawn of targeted bacteria and incubated aerobically, however, when the above assay was repeated under anaerobic conditions *Pcb*1692 lost the ability to produce carbapenem but not bacteriocins. Taken together, these findings suggest that *Pcb*1692 deploys carbapenem and bacteriocins to inhibit growth of targeted bacteria either in the rhizosphere, phyllosphere or on the surface of potato tubers. Once inside potato tubers, the T6SS of *Pcb*1692 is activated and alongside its arsenal of bacteriocins, inhibits growth of targeted bacteria. This hypothesis is supported by our transcriptomic data which demonstrated that while genes associated with the *Pcb*1692 T6SS and bacteriocin production were massively upregulated *in planta*, genes associated with carbapenem biosynthesis were not differentially expressed at 24 and 72 hours post-inoculation (47).

It is important to note that because of the difficulty associated with identification and quantification of antibiotics and bacteriocins produced by bacteria *in planta*, most studies, including the current study, make inferences based on *in vitro* detection of these compounds and extrapolations/inferences are made to *in planta* conditions based on the assay conditions which mimic plant apoplast (minimal media and anaerobic conditions) (65). Against this background it is possible that *Pcb*1692 does not produce carbapenem *in planta* but is able to produce bacteriocins. Although bacteriocin production is usually induced *in vitro* by DNA damaging agents, which trigger SOS response leading to the production of bacteriocins (10, 14), it is possible that bacteria can produce bacteriocins *in planta*. In support of this hypothesis, recent studies have shown that bacterial SOS can be triggered directly or indirectly by different factors such as nutrient starvation, iron deficiency, chromate stress, acidic and alkaline stress, oxidative stress and β-lactam antibiotics (10, 66-69). All the above scenarios can be encountered by *Pcb in planta*, for example, one of the first stresses encountered by *Pcb*, SREs and other endophytes in potato tubers is acidic pH, which SREs must neutralize to be able to produce PCWDEs (65, 70, 71). Similarly, intake of excess iron can lead to oxidative burst in bacteria through Fenton reaction (72). In addition, the presence of chromate in potato tubers (45, 73) and production of β-lactam antibiotics by either bacteria endophytes or the host plant can trigger an SOS response and hence bacteriocin production (74, 75)

Previous studies have shown that carbapenem production by *Pcc* and *Serratia* is regulated by quorum sensing and SlyA (26). The current working model on carbapenem production is that at a low cell density, when AHL concentration is low, CarR cannot perform AHL-dependent transcription of the *car* genes from the *car*A promoter (26, 76). However, once quorum is reached and AHL molecules are available, the formation of CarR-AHL complex enables both self-activation of *car*R, and activation of the downstream *car* genes through the *carA* promoter region (77). Conversely, regulation of carbapenem production by SlyA is independent of AHL molecules and the putative SlyA binding site in the *car* gene cluster is unknown (76). The above model is consistent and applicable to *Pcb*1692 because our results demonstrate that while wildtype *Pcb*1692 produced carbapenem on LB the isogenic mutant strains, *Pcb*1692Δ*exp*I and *Pcb*1692Δ*sly*A do not produce detectable levels of carbapenem, under the same experimental conditions. These findings are also supported by our *in silico* analyses (Fig 4) which show a highly conserved genetic organisation of the *car* gene cluster in several strains of *Pcb* including *Pcb*1692 and several strains of *Pcc*. Hence, the conservation of similar mechanisms of carbapenem regulation in these bacteria is presumable.

The same analysis also showed that publicly available genome sequences of several strains of *Pcb, Pcc, P. parmentieri*, and *P. wasabiae* do not have genes associated with carbapenem biosynthesis, however, some of these strains have gene homologs of the *car*F and *car*G genes encoding immunity proteins for carbapenem, suggesting they could resist the carbapenem produced by closely related species. However, the exact role of CarF and CarG in carbapenem resistance is not well understood because 1) bacteria produce different isoforms of carbapenem (51), and 2) carbapenem resistance has been associated with different class C family of β-lactamases produced by several bacteria (30). Furthermore, *car*F and *car*G can be transcribed from the *car*A promoter sequence by CarR in a quorum dependent manner, however, these genes can also be weakly transcribed from a promoter located within *car*G (76). Therefore, the absence of these promoter sequences in SREs with a truncated *car* gene clusters suggests that *car*F and *car*G may not be transcribed in these bacteria (Fig 4). In support of this hypothesis, this study showed that environmental strains of *Pcb* (XT3, XT10, G4P5 and G4P7) were all inhibited by the carbapenem produced by *Pcb*1692 alongside other *Pcb* strains. PCR amplification failed to amplify the *car*F and *car*G genes in *Pcb* (XT3, XT10, G4P5 and G4P7) (results not shown), however, because of the possibility of sequence variation between closely related bacterial strains, one cannot rule out the presence or absence of a gene in a given bacterial genome solely based of PCR. Furthermore, because of the genetic heterogeneity of *Pcb* strains and SREs in general, as seen by the recent taxonomic reclassification of several SREs (34, 78), it is possible that the genome sequences of *Pcb* strains XT3, XT10, G4P5 and G4P7 do not encode homologs of CarF and CarG, or do not encode the cognate CarF and CarG homologs providing resistance to carbapenem produced by *Pcb*1692. Alternatively, if present, the *carF* and *G* genes of *Pcb* strains XT3, XT10, G4P5 and G4P7may not be transcribed if these strains have a truncated *car* gene cluster similar the ones described in our comparative analysis.

Furthermore, while gene homologues associated with the *car* gene cluster could not be identified in the genome sequences of most *Dickeya* spp. whose genome sequences are publicly available, the *car* gene cluster was present in one strain of *D. dadantii* and *D. zeae* alongside three strains of *D. chrysanthemi* (Fig 4). Interestingly, in *Dickeya* spp we observed a different genetic organisation of the *car* gene cluster and the replacement of *car*R with a LacI family of transcriptional regulators (Fig 4). These findings raise the possibility that the *Dickeya* LacI may respond to environmental or intracellular cues which are different from those associated with CarR-mediated transcriptional regulation of the *car* gene cluster in *Pcc* and *Pcb*1692. However, it remains to be determined if these *Dickeya* spp produce carbapenem, under what conditions and the role played by AHL and SlyA in the regulation of the *car* gene cluster in these bacteria.

Iron is an essential element which acts as a cofactor of several enzymes, a constituent of heme and is required for biosynthesis of iron-sulphur clusters (Fe-S) found in several proteins (79, 80). The limited amount of iron found in plant apoplast suggests that bacteria have to compete to obtain this scarce metal, with some bacteria producing species-specific siderophores which typifies exploitative competition (16, 81). Despite its involvement in several biological processes, excess iron is toxic to bacteria and intracellular levels of iron are regulated by the ferric uptake regulator (Fur) (82). In the presence of iron, Fur binds to iron and the Fur^Fe2+^ complex represses expression of several genes including iron uptake gene while concomitantly activating expression of genes associated with iron efflux, virulence, stress response and several metabolic process (79, 80, 83-86). Fur regulates genes by either directly binding to their promoter sequences or indirectly by recruiting small regulatory RNA which then regulate gene expression (82, 84, 85, 87, 88). The Fur regulon has been described for several bacteria including genes regulated by Fur under iron limiting or excess iron conditions (68, 79, 80, 82, 89, 90). Interestingly, our results also showed that carbapenem production by *Pcb*1692 is dependent on iron and regulated by Fur. This is based on the fact that *Pcb*1692Δ*fur* mutant strain lost the ability to produce carbapenem compared to the wild-type strain. Similarly, the results showed that wild-type *Pcb*1692 only produced carbapenem when M9 was supplemented with iron. Identification of putative Fur binding sites in *sly*A and *car*R promoter regions in *Pcb*1692, *Pcc, Pco* and *Pca* further suggests that the iron-dependent regulation of carbapenem by Fur-Fe^2+^ through activation of *slyA* and *carR* may be conserved across the *Pectobacterium* genus. In addition, the results also showed that *Pcb*1692 produces carbapenem on LB while the isogenic mutants of *sly*A, *fur, exp*I do not, suggesting that the amount of iron in yeast extracts found in LB may be enough for transcriptional activation of the *Pcb*1692 *car* genes.

The requirement for iron and Fur in the biosynthesis of carbapenem is consistent with the finding that CarE is a putative ferredoxin with 2 Fe-S clusters while the carbapenem synthase CarC protein is an oxygenase which requires iron and oxygen and 2-oxoglutarate as co-factors (51, 91). Previous studies have shown that under iron limiting conditions, Fur upregulates transcription of genes encoding essential iron-containing proteins including the small regulator RNA, RhyB (68, 87, 92). RhyB in turn degrades mRNA of genes encoding non-essential iron-containing proteins in what has been described as an iron-sparing response suggesting that CarC and CarE may be post-transcriptionally repressed under iron-limiting conditions (82). The requirement for oxygen by CarC is also supported by our findings which showed that that *Pcb*1692 does not produce carbapenem under anaerobic conditions. Furthermore, in *Pectobacterium* spp. *exp*R which is convergently transcribed from *exp*I is the transcriptional activator of RmsA, which is a global repressor of several genes including SlyA and ExpI (93-96). In addition, RmsA production is repressed by AHL while deletion of *exp*R leads to increased production of AHL molecules (93, 97). Our *in silico* analysis identified a putative *fur* binding site in the promoter sequence of *exp*R suggesting that in the presence of iron, Fur^Fe2+^ might repress expression of *exp*R and *rms*A leading to increased production of SlyA and AHLs which are both needed for transcription of *car* genes. Conversely, in the absence of iron, ExpR activates production of RmsA which degrades mRNA of SlyA hence no carbapenem production.

We hypothesize that in LB medium, in which bacterial cells grow relatively fast thus promoting accumulation of AHL, Fur-Fe^2+^ binds to the promoter sequences of *car*R and *sly*A thereby activating transcription of these regulators leading to carbapenem production. However, in minimum medium (absence of iron) slow growing bacteria produce limited amounts of AHL molecules, and Fur is unable to activate transcription of *car*R and *sly*A hence carbapenem production is impaired. Conversely, we observed a slight increase in the growth rate of *Pcb*1692 when minimum media was supplemented with iron, which could translate into increased production of AHL molecules. Under these conditions, Fur-Fe^2+^ promotes transcription of *car*R and *sly*A and CarR-AHL and SlyA together mediates transcription of the *car* genes. In summary, the requirement for iron by both CarC and CarE is consistent with the observed iron-dependent regulation of the *car* gene cluster by Fur in *Pcb*1692. Additionally, coupling the iron monitoring/uptake machineries with the transcriptional regulation of carbapenem may be a conserved trait in other *Pectobacterium* spp. as suggested by the binding sites detection.

In conclusion, our results show that *Pcb* encounters several different endophytes in potato tubers while some of the interactions may be cooperative, other interactions are antagonistic. Competition by interference is mediated by recruitment of *Pcb*1692 T6SS *in planta* possibly on cue to as yet unknown plant signals. Through this contact dependant mechanism, *Pcb* is used to inhibit growth of several bacteria. Furthermore, *Pcb*1692 produces carbapenem and bacteriocins, which inhibit several gram-negative and gram-positive bacteria including *Bacillus subtilis* and *B. cereus*. While the *Pcb*1692 can produce bacteriocins aerobically and anaerobically, carbapenem production was found to be dependent on availability of oxygen. Finally, the results showed that carbapenem production is *Pcb*1692 regulation by the concerted action of Fur, SlyA and ExpI.

## Materials and methods

### Strains and growth conditions

All bacterial strains used in this study are listed in Table 1. Bacterial strains were cultured in either Luria-Bertani (LB) medium or M9 minimum medium (98) supplemented with 0.4% sugar (glucose, sucrose or glycerol) and grown aerobically at 37°C or 28°C with or without agitation, as required by the given experimental conditions. Growth medium was supplemented with either 100 μg/ml ampicillin, 50 μg/ml kanamycin, 15 μg/ml gentamycin, 15 μg/ml tetracycline or 50 μg/ml chloramphenicol. Antibiotics were purchased from (Sigma-Aldrich).

**Table 1.** List of bacteria used in bacterial competition. **Table1-List of bacterial strains used in this study:** UK = United Kingdom, USA = United States of America, FABI = Forestry and Agricultural Biotechnology Instituted, University of Pretoria. Bacterial strains whose genome sequence has been determined are indicated as “sequenced” in the Description column. Bacteria type strains and indicated by the by a superscript T (^T^) following the bacterial name. *P* = *Pectobacterium, Pcc* = *Pectobacterium carotovorum* subsp. *carotovorum*

| Bacterial strains | Description (Host and country of isolation) | Sources |
| --- | --- | --- |
| *Pectobacterium carotovorum* subsp. *brasiliense* (*Pcb*) |  |  |
| *Pcb* 1692 | Potato, Brazil, sequenced | (99) |
| *Pcb* G4P5 | Potato, South Africa | FABI |
| *Pcb* G4P7 | Potato , South Africa | FABI |
| *Pcb* XT3 | Potato , South Africa | FABI |
| *Pcb* XT10 | Potato , South Africa | FABI |
| *Pcb* 358 | Potato, South Africa | FABI |
| *Pcb* CC1 | Cucumber, South Africa | FABI |
| *Pcb* CC2 | Cucumber, South Africa | FABI |
| *Pcb* HPI01 | Cucumber, South Africa, sequenced | (100) |
| *Pcc*1 | Unknown, South Africa | FABI |
| *Pcc*2 | Unknown, South Africa | FABI |
| *PccBR1* | Unknown, South Africa | FABI |
| *PccBR3* | Unknown, South Africa | FABI |
| *PccBR7* | Unknown, South Africa | FABI |
| *PccRP6P2* | Unknown, South Africa | FABI |
| *PccGP6P1* | Unknown, South Africa | FABI |
| *PccYP10R2* | Unknown, South Africa | FABI |
| *PccRP17P1* | Unknown, South Africa | FABI |
| *PccRP17P2* | Unknown, South Africa | FABI |
| *P. atrosepticum* ATCC 33260<sup>T</sup> | Potato, UK, sequenced | (101) |
| *P. carotovorum* ATCC 15713<sup>T</sup> | Potato, sequenced | (102) |
| *P. wasabiae* ATCC 43316<sup>T</sup> | *Eutrema wasabi* ,Japan, sequenced | (103) |
| *P. cypripedii* PDDCC 1591<sup>T</sup> | *Cypripedium sp.*,USA | (104) |
| *Dickeya paradisiaca* ATCC 33242<sup>T</sup> | *Musa paradisiaca* var. *dominico*,<br>Colombia, sequenced | (105) |
| *P. betavascularum* LMG 2398 | Potato, Romania | FABI |
| *P. carotovorum* subsp. *odoriferum* | South Africa | FABI |
| *Salmonella typhimurium* | South Africa` | FABI |
| *Escherichia coli* | South Africa | FABI |
| *Dickeya dadantii* LMG 25991<sup>T</sup> | *Pelargonium capitatum*, Comoros,<br>sequenced | (106) |
| *Erwinia chrysanthemi* ATCC 11663<sup>T</sup> | *Chrysanthemum* , USA, sequenced | (107) |
| *Erwinia rhapontici* LMG 2645 | Pear, UK | FABI |
| *Serratia marcescens subsp marcescens*<br>ATCC 13880<sup>T</sup> | Pond water, Czech Republic, sequenced | (108) |
| *Serratia fiscaria* ATCC 33105<sup>T</sup> | Fig, | (109) |
| *Bacillus subtilis* | Unknown | FABI |
| *Bacillus cereus* | Unknown | FABI |
| *Pantoea ananatis* LMG2665<sup>T</sup> | Onion, Brazil, sequenced | (110) |
| *Pantoea stewartii subsp. indologenes*<br>ATCC 51785<sup>T</sup> | *Setaria italic*, India, sequenced | (111) |
| *Enterobacter cowanii* | *Eucalyptus grandis*, Uruguay | FABI |

### DNA isolation, Miseq Illumina sequencing and data analysis

Six diseased (soft rot disease) and six healthy potato tuber were collected from Gauteng, North West and Limpopo province in South Africa. Genomic DNA (gDNA) was extracted from 1g of fresh tissue using the ZYMO RESEARCH Quick-gDNA™ MiniPrep kit, according to the manufacturer’s instruction. DNA from either healthy or diseased tubers was pooled and aliquots used for 16S rRNA PCR amplification and Illumina Miseq sequencing. The 16S rRNA PCR was performed to amplify v3-v4 variable regions using primers 419F (ACTCCTACGGGAGGCAGCAG) and 806R (GGACTACHVGGGTWTCTAAT) (112). The PCR reaction mixture consisted of 2.5 mM dNTPs, 10× DreamTaq buffer (supplemented with 20 mM MgCl_2_), 0.5 U DreamTaq Polymerase (Fermentas), 10 μM each forward and reverse primer, 100 ng DNA template and nuclease free water up to the final reaction volume of 25 µl. PCR amplification conditions were as follows: denaturation at 95°C for 3 min, followed by 35 cycles at 95°C 30s, 58°C 45s, 72°C 2 min and the final extension at 72°C for 5 min. The purified PCR products (260/280 ratio ≥ 1.8) were sent to the Agricultural Research Council (ARC) Biotechnology platform, Pretoria, South Africa for 16S metagenomics library preparation, modified with Illumina specific adapters (113). Thereafter, the pooled samples were subjected to Miseq Illumina sequencing. Paired-end reads were trimmed using Trimmomatic v0.38 (114), allowing minimum nucleotide quality average of 30 per 5 bp window in the sequences. Illumina adapters were removed using Cutadapt v1.18 (115). Chloroplast sequences were filtered out based on sequence identity assessed through Blastn (116) with 16S rRNA acquired from *Solanum tuberosum*. High-quality reads were then submitted to SILVAngs rRNA-based data analysis pipeline under default settings (43). Next, the Krona package (117) was used to generate radial visualization of the resulting taxonomic profiles. The 16S metagenomics raw data has been deposited in The Sequence Read Archive database on NCBI under the accession number PRJNA509544.

### Plant extracts

Potato tuber extracts were prepared as previously described, with slight modifications (118). In summary, five grams of surface sterilized potato tubers (flesh and peel) were ground in 100ml of distilled water using a blender. Plant tissue was removed by repeated centrifugation at 5000 rpm for 30mins at 4°C. The clarified plant extract was filter sterilized twice using a 0.2µm filters, and aliquots stored at -16°C. Iron was selectively removed from potato tuber extracts by adding 200µM of 2,2’-dipyridyl (Sigma Aldrich) dissolved in chloroform in a 1:1 ratio. The resulting mixture was vortexed for 2mins and iron free supernatant collected by centrifugation at 5000rpm for 10mins at 4°C. The M9 agar plates were then supplemented with a 1:10 dilution of either plant extracts or plant extracts pre-treated with 2,2’-dipyridyl.

### Orthology predictions and binding sites analyses

Orthologs of bacteriocins and *car* genes were predicted by using OrthoMCL (119) pipeline to analyse complete protein datasets from 100 *Pectobacterium* and *Dickeya* strains obtained from the RefSeq database as previously described (47). Upstream regions of *expR, sly*A, *car*R and *car*A from *Pectobacterium betavasculorum* strain NCPPB2795, *Pectobacterium atrosepticum* strain ICMP19972 *carotovorum* subsp. *carotovorum* strain NCPPB312, and *Pcb*1692 genomes were retrieved from the NCBI online database (www.ncbi.nlm.nih.gov). Then the respective sequences of those regions were scanned using FIMO software from MEME package (49) for the presence of putative Fur binding sites. This analysis was conducted with support of all known Fur binding sites found in MEME databases for prokaryotes: Prodoric (release 8.9) (120), RegTransBase v4 (121), and CollectTF (122).

### *In vitro* bacterial competition assays

Inter and intra species bacterial competition assays were performed as previously described (25). The complete list of bacteria used in this assay is provided in Table 1. In summary, targeted bacteria were transformed with plasmid pMP7605 conferring gentamycin resistance (123). Overnight cultures of *Pcb*1692 and targeted bacteria were normalized to an OD_600_ = 0.1, mixed in a 1:1 ratio and 20µl spotted on LB or M9 agar containing no antibiotics. When required, M9 was supplemented with 0.01, 1, 10 or 50 µM of ferric sulphate (FeSO_4_.7H_2_O), magnesium sulphate (MgSO_4_.7H_2_O), manganese chloride (MnCl_2_.4H_2_O), zinc sulphate (ZnSO_4_.7H_2_O), nickel sulphate (NiSO_4_), cobalt chloride (CoCl_2_.H_2_O) or copper chloride (CuCl.H_2_O).

Bacterial spots were briefly air dried and allowed to grow for 16 hours at 28°C. When required, bacterial competition assays were performed anaerobically by placing agar plated in an anaerobic jar (Merk) containing moistened Anaerocult^®^ A (Merk) which absorbs oxygen and a moistened blue Anaerotest^®^ strip (Merk) which turns white in the absence of oxygen. Anaerobic jars were then tightly sealed and incubated overnight at 4°C. Thereafter, overnight spots were scraped off, serially diluted and plated on LB agar supplemented with gentamycin (15µg/ml). The effect of iron on bacterial competition was determined using the following agar plates: 1) LB agar, 2) M9 minimal agar, 3) M9 supplemented with FeSO_4_.7H_2_O (10 µM), 4) M9 supplemented with potato tuber extracts, 5) M9 supplemented with FeSO_4_.7H_2_O (10 µM) and potato tuber extracts, 6) M9 supplemented with potato tuber extracts pre-treated with the iron chelator 2,2’-dipyridyl (200µM) and 7) complementation of plate 6 listed above by the addition FeSO_4_.7H_2_O (10µM) to potato tubers pre-treated with the 2,2’-dipyridyl (200µM). All assays were repeated three times and results presented as CFU/ml of target bacteria.

### Generation of *Pectobacterium carotovorum* subsp. *brasiliense* mutant strains

The different *Pcb*1692 mutant strains were generated by site-directed mutagenesis using the lambda red recombination technique (124). In summary, overlap extension polymerase chain reaction (PCR) was used to generate a gene knockout cassette by fusing the upstream and downstream regions flanking the targeted gene to the kanamycin resistance genes (25, 125). The fused PCR product was then electroporated into *Pcb*1692 harbouring pKD20 and transformants were selected on nutrient agar supplemented with either 50 µg/ml kanamycin or 50 µg/ml chloramphenicol. The list of primers used in this study is provided in Table 2. The integrity of each *Pcb*1692 mutant strain was confirmed by PCR analyses, nucleotide sequencing and Southern blot analysis (results not shown). The list of mutant strains generated in this study are provided in Table 3.

**Table 2.**
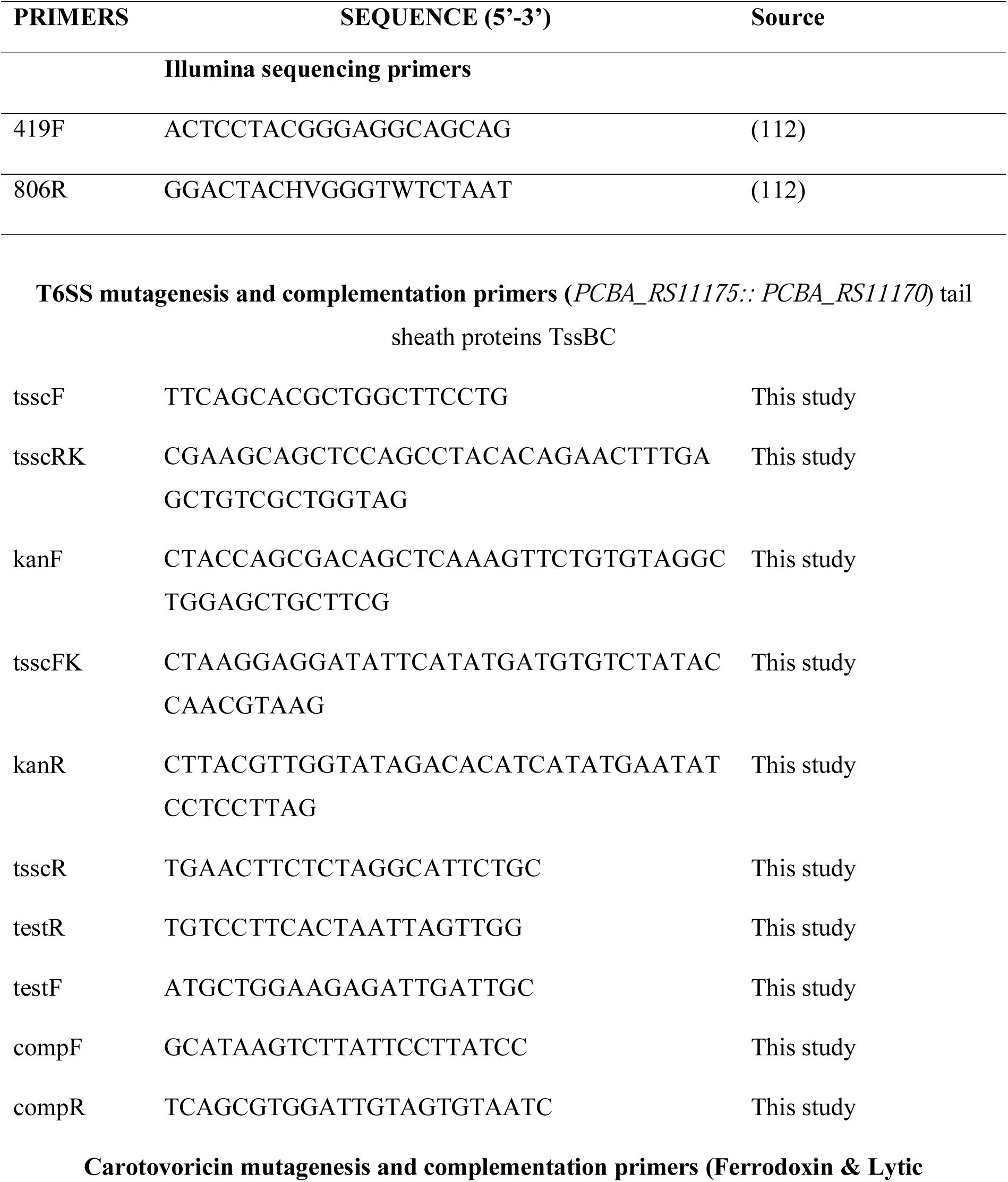
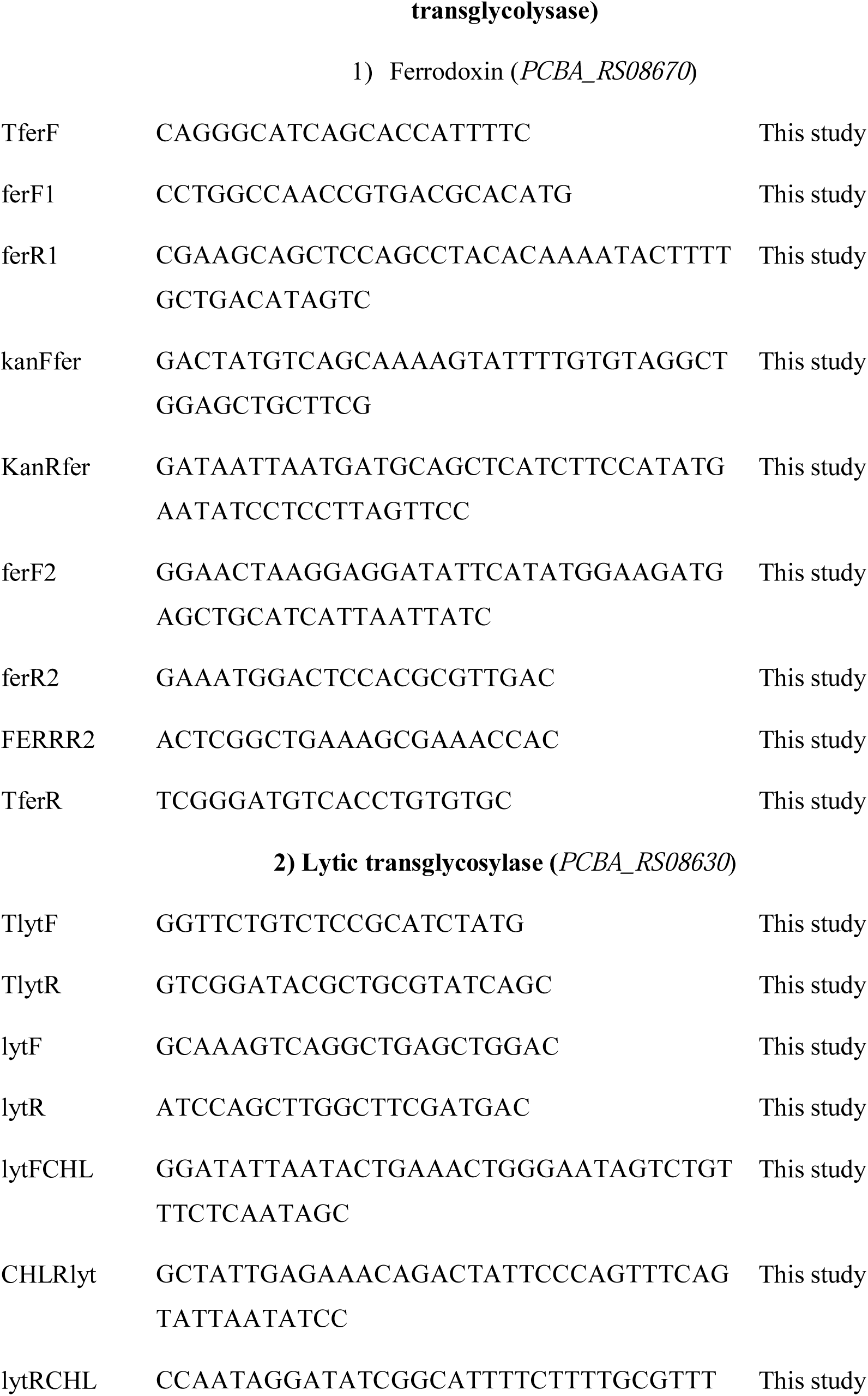
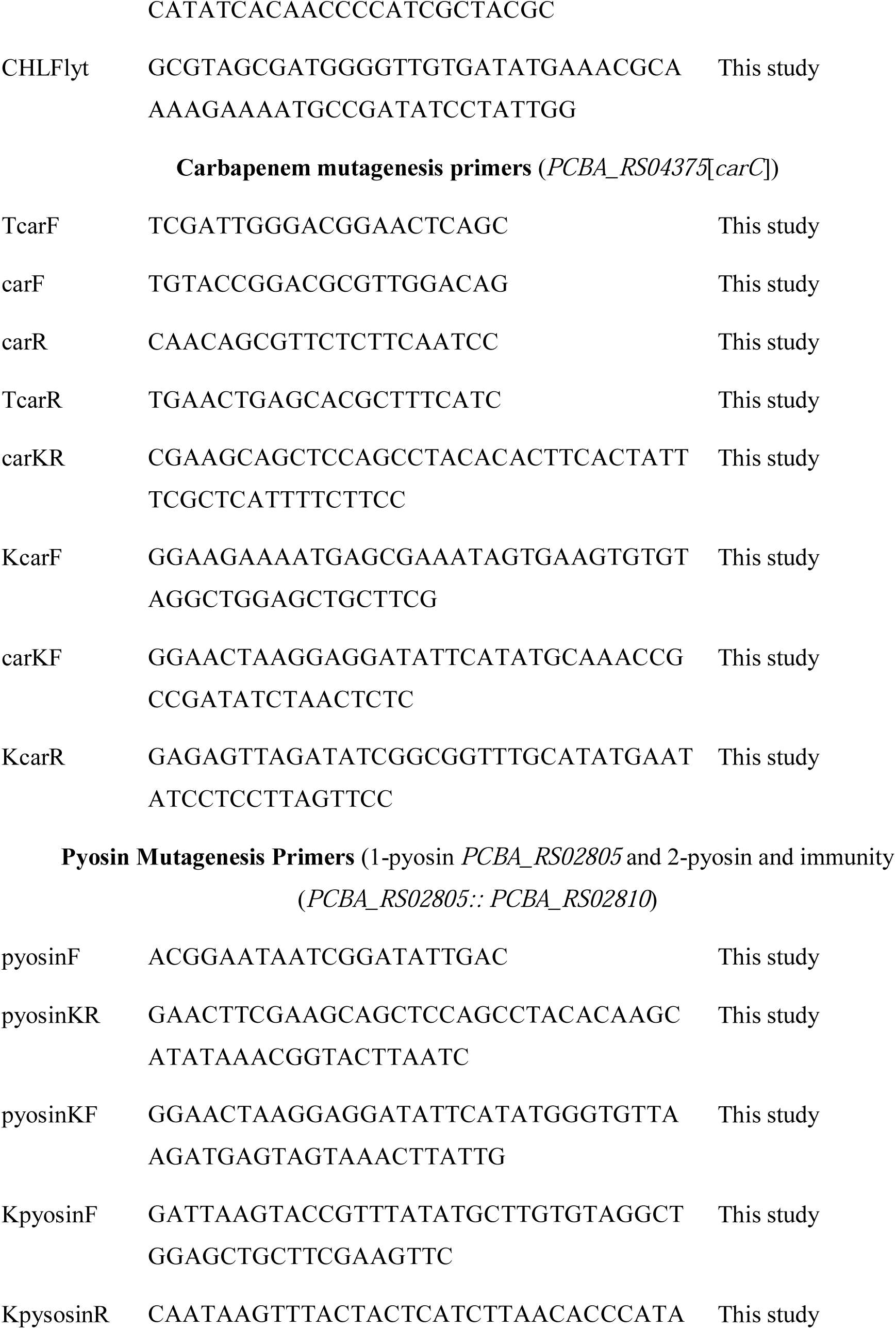
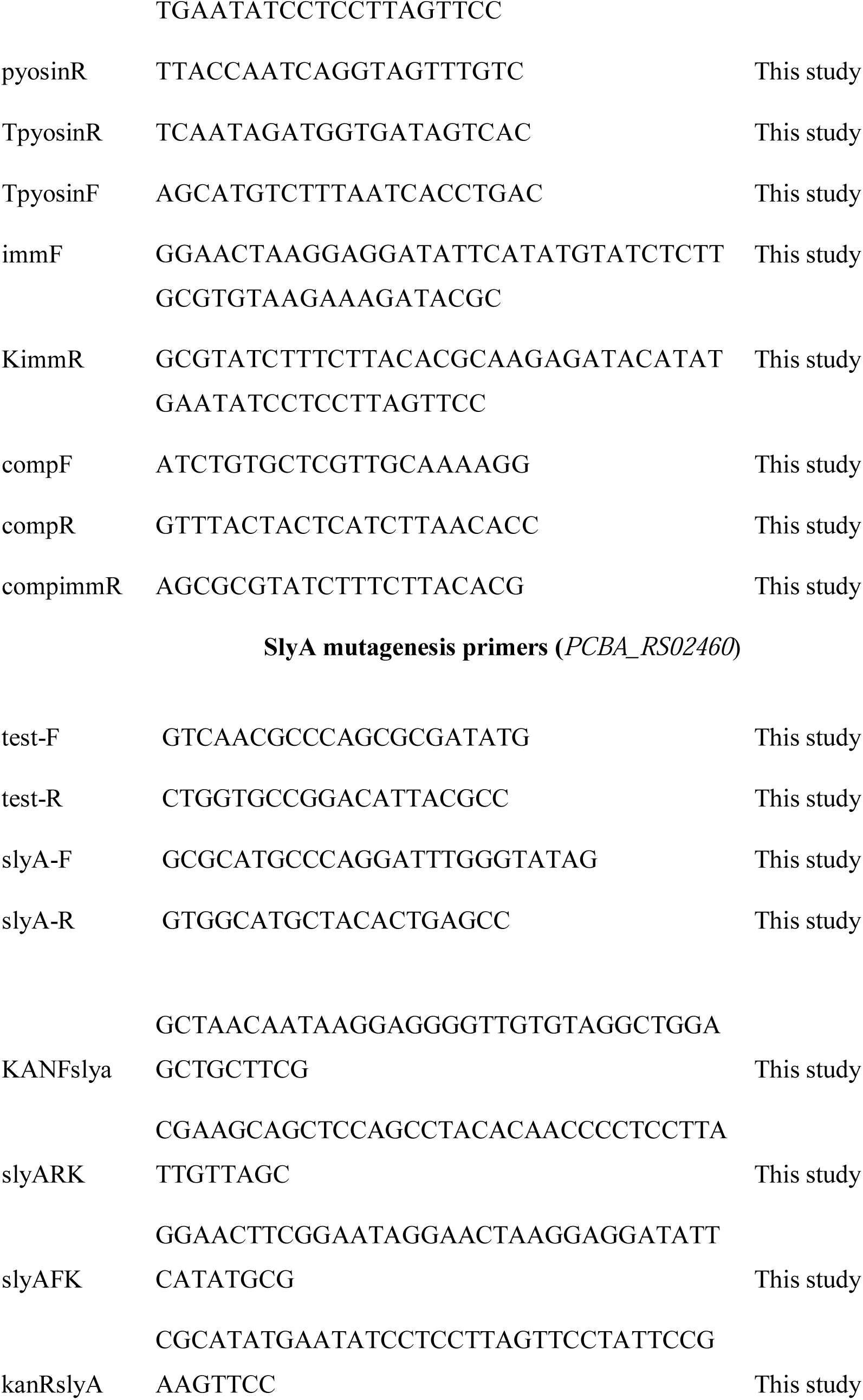
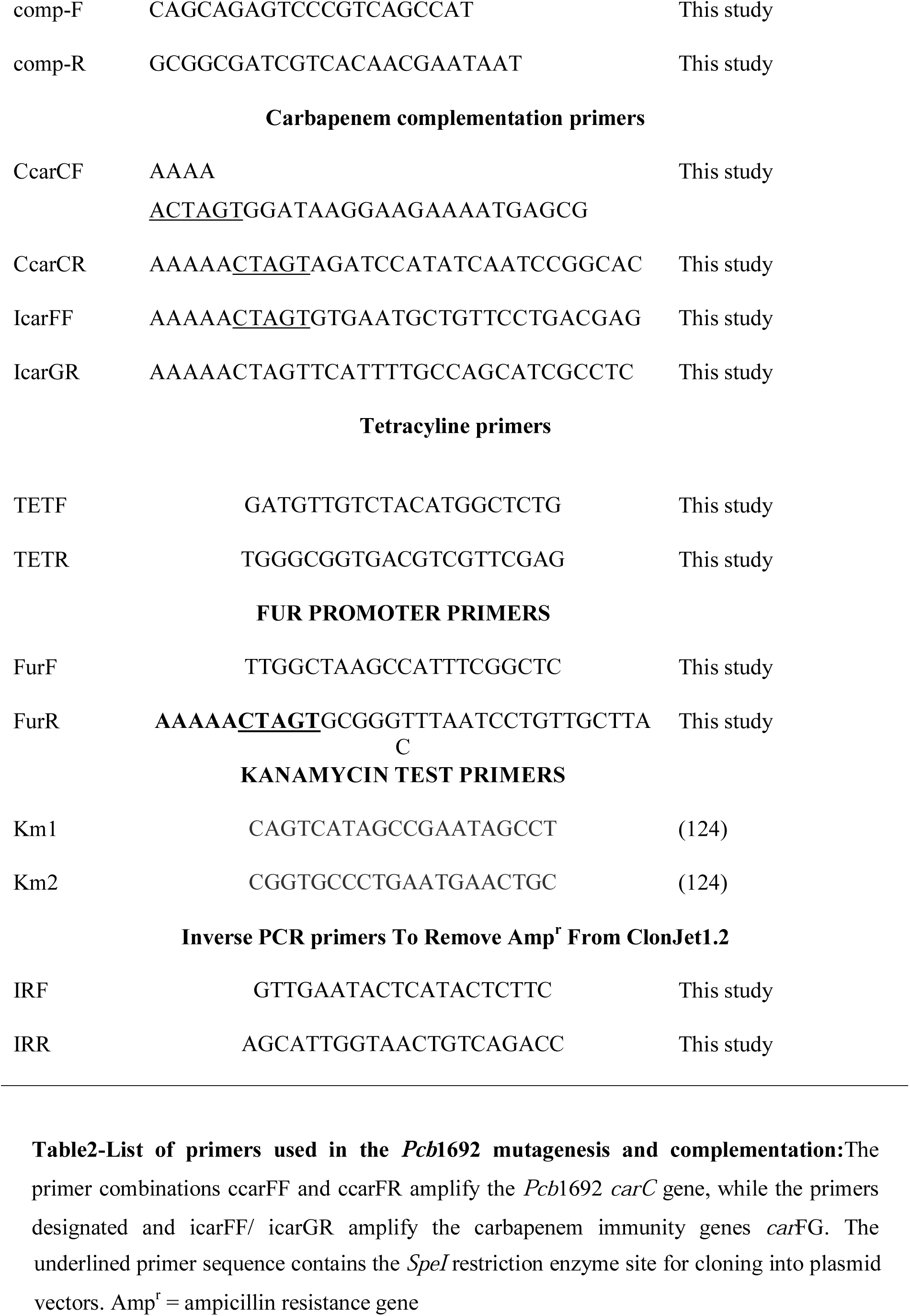
List of primers used in the *Pcb*1692 mutagenesis and complementation. The primer combinations ccarFF and ccarFR amplify the *Pcb*1692 *carC* gene, while the primers designated and icarFF/ icarGR amplify the carbapenem immunity genes *car*FG. The underlined primer sequence contains the *SpeI* restriction enzyme site for cloning into plasmid vectors. Amp^r^ = ampicillin resistance gene

**Table 3.**
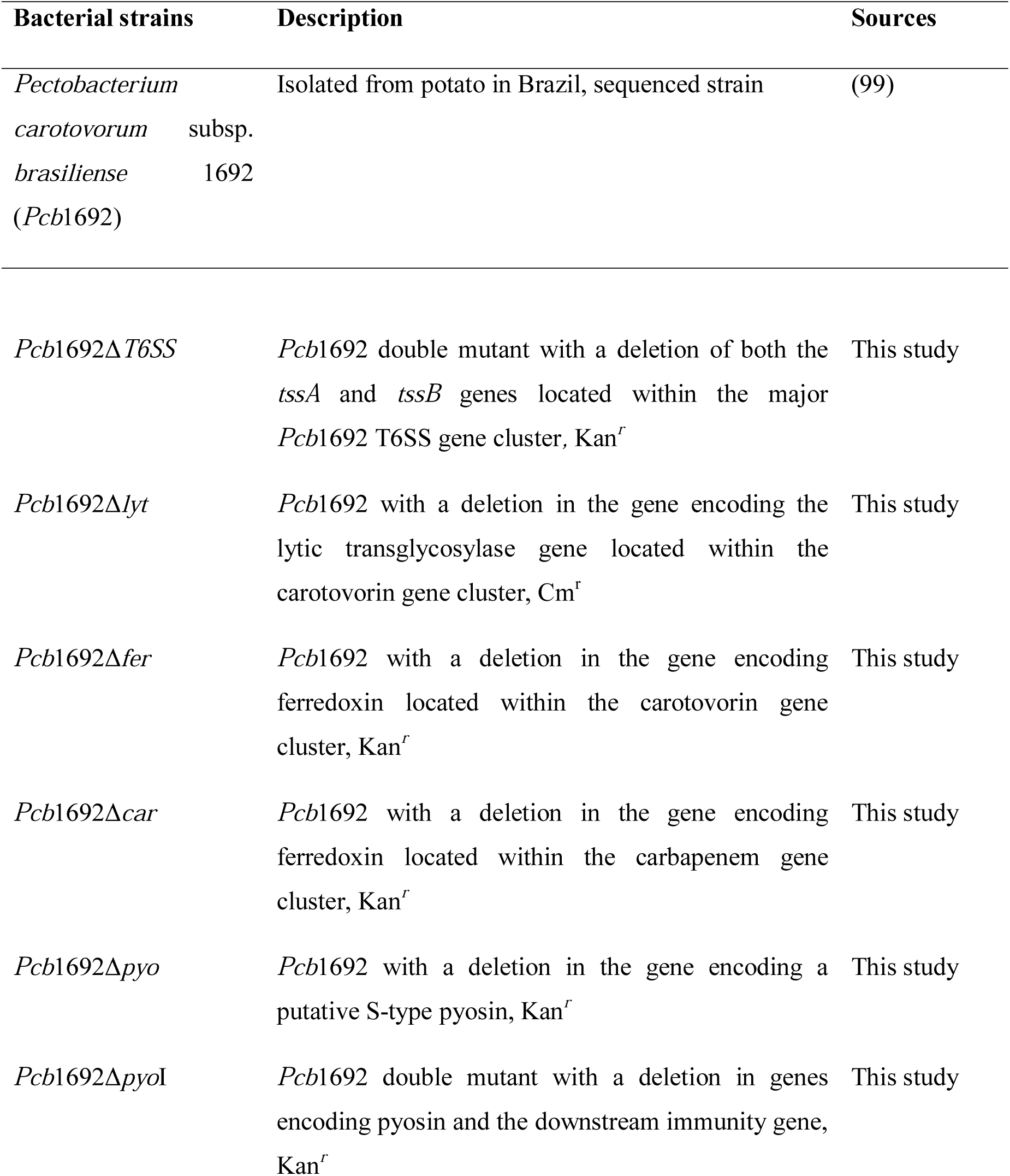
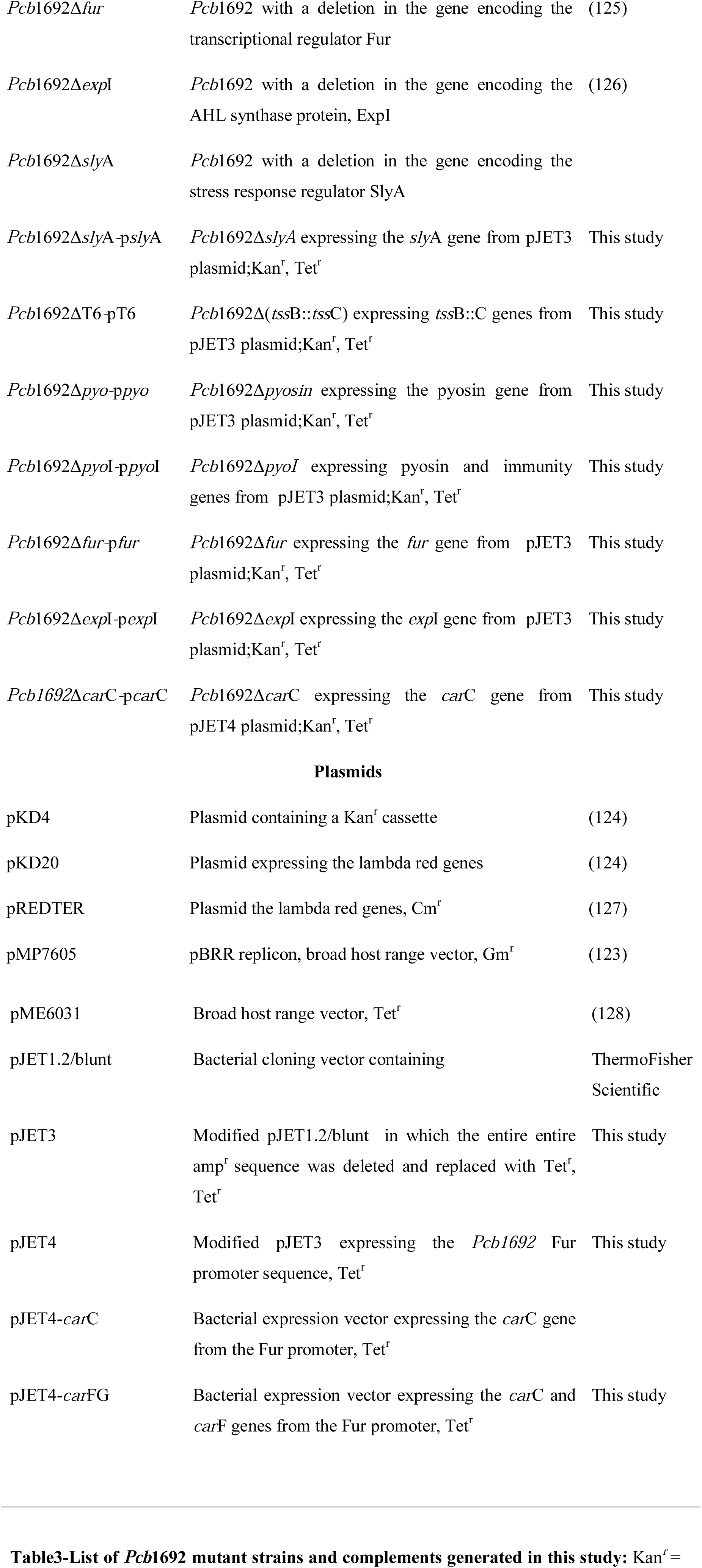
List of *Pcb*1692 mutant strains and complements generated in this study. Kan^*r*^ = kanamycin resistance, Tet^r^ = tetracycline resistance, Cm^r^ = chloramphenicol resistance.

| Bacterial strains | Description | Sources |
| --- | --- | --- |
| *Pectobacterium carotovorum* subsp. *brasiliense* 1692 (*Pcb1692*) | Isolated from potato in Brazil, sequenced strain | (99) |
| *Pcb1692*Δ*T6SS* | *Pcb1692* double mutant with a deletion of both the *tssA* and *tssB* genes located within the major *Pcb1692* T6SS gene cluster, Kan<sup>r</sup> | This study |
| *Pcb1692*Δ*lyt* | *Pcb1692* with a deletion in the gene encoding the lytic transglycosylase gene located within the carotovarin gene cluster, Cm<sup>r</sup> | This study |
| *Pcb1692*Δ*fer* | *Pcb1692* with a deletion in the gene encoding ferredoxin located within the carotovarin gene cluster, Kan<sup>r</sup> | This study |
| *Pcb1692*Δ*car* | *Pcb1692* with a deletion in the gene encoding ferredoxin located within the carbapenem gene cluster, Kan<sup>r</sup> | This study |
| *Pcb1692*Δ*pyo* | *Pcb1692* with a deletion in the gene encoding a putative S-type pyosin, Kan<sup>r</sup> | This study |
| *Pcb1692*Δ*pyoI* | *Pcb1692* double mutant with a deletion in genes encoding pyosin and the downstream immunity gene, Kan<sup>r</sup> | This study |
| *Pcb1692Δfur* | *Pcb1692* with a deletion in the gene encoding the transcriptional regulator Fur | (125) |
| *Pcb1692ΔexpI* | *Pcb1692* with a deletion in the gene encoding the AHL synthase protein, ExpI | (126) |
| *Pcb1692ΔslyA* | *Pcb1692* with a deletion in the gene encoding the stress response regulator SlyA |  |
| *Pcb1692ΔslyA-pslyA* | *Pcb1692ΔslyA* expressing the *slyA* gene from pJET3 plasmid; Kan<sup>r</sup>, Tet<sup>r</sup> | This study |
| *Pcb1692ΔT6-pT6* | *Pcb1692Δ(tssB::tssC)* expressing *tssB::C* genes from pJET3 plasmid; Kan<sup>r</sup>, Tet<sup>r</sup> | This study |
| *Pcb1692Δpyo-ppyO* | *Pcb1692Δpyosin* expressing the pyosin gene from pJET3 plasmid; Kan<sup>r</sup>, Tet<sup>r</sup> | This study |
| *Pcb1692ΔpyoI-ppyOI* | *Pcb1692ΔpyoI* expressing pyosin and immunity genes from pJET3 plasmid; Kan<sup>r</sup>, Tet<sup>r</sup> | This study |
| *Pcb1692Δfur-pfur* | *Pcb1692Δfur* expressing the *fur* gene from pJET3 plasmid; Kan<sup>r</sup>, Tet<sup>r</sup> | This study |
| *Pcb1692ΔexpI-pexpI* | *Pcb1692ΔexpI* expressing the *expI* gene from pJET3 plasmid; Kan<sup>r</sup>, Tet<sup>r</sup> | This study |
| *Pcb1692ΔcarC-pcarC* | *Pcb1692ΔcarC* expressing the *carC* gene from pJET4 plasmid; Kan<sup>r</sup>, Tet<sup>r</sup> | This study |
| Plasmids |  |  |
| pKD4 | Plasmid containing a Kan<sup>r</sup> cassette | (124) |
| pKD20 | Plasmid expressing the lambda red genes | (124) |
| pREDTER | Plasmid the lambda red genes, Cm<sup>r</sup> | (127) |
| pMP7605 | pBRR replicon, broad host range vector, Gm<sup>r</sup> | (123) |
| pME6031 | Broad host range vector, Tet<sup>r</sup> | (128) |
| pJET1.2/blunt | Bacterial cloning vector containing | ThermoFisher Scientific |
| pJET3 | Modified pJET1.2/blunt in which the entire entire amp<sup>r</sup> sequence was deleted and replaced with Tet<sup>r</sup>, Tet<sup>r</sup> | This study |
| pJET4 | Modified pJET3 expressing the *Pcb1692* Fur promoter sequence, Tet<sup>r</sup> | This study |
| pJET4-*carC* | Bacterial expression vector expressing the *carC* gene from the Fur promoter, Tet<sup>r</sup> |  |
| pJET4-*carFG* | Bacterial expression vector expressing the *carC* and *carF* genes from the Fur promoter, Tet<sup>r</sup> | This study |

### Complementation of the *Pcb*1692 mutant strains

The list of primers used in this study is provided in Table 2. The *Pcb*1692 *fur, sly*A, *exp*I, *PCBA_RS02805* (pyocin, designated *pyo*), *PCBA_RS02805::PCBA_RS02810* (pyocin and immunity genes, designated *pyo*I) and the T6SS genes *tss*A/B including their putative promoter sequences were PCR amplified and individually cloned into CloneJet 1.3 to generate p*fur*, p*sly*A, p*exp*I, p*pyo*, p*pyo*I and pT6, respectively. The tetracycline resistance gene (tet^r^) with its downstream promoter sequence was amplified from plasmid pME6031 (128) with primers tetF/tetR. The ampicillin resistance gene located in plasmids p*fur*, p*sly*A, p*exp*I, p*pyo*, p*pyo*I and pT6 was removed by inverse PCR, using primers IrF/IrR. The linearized vectors were then ligated to tet^r^ to generate pJET3-*fur*, pJET3-*sly*A and pJET3-*exp*I, pJET3-*pyo*, pJET3-*pyo*I and pJET3-T6. The *Pcb*1692 *fur* promoter sequence was amplified with FurF/R and ligated to cloneJet 1.3 to generate p*fur-*1. The amp^r^ gene in p*fur-*1 was replaced with tet^r^, as previously described, generating pJET4. The promoterless *Pcb*1692 *car*C and *car*FG genes were amplified by PCR, followed by restriction digested with SpeI and cloned into pJET4 previously digested with SpeI, to generate pJET4-*car*C and pJET4-*car*FG, respectively. Plasmids pJET3-*fur*, pJET3-*sly*A, pJET3-*exp*I, pJET4-*car*C, pJET3-*pyo*, pJET3-*pyoI* and pJET3-T6 were transformed into their corresponding mutant strains to generate the complemented mutant strains, *Pcb*1692Δ*fur-*p*fur, Pcb*1692Δ*sly-*p*sly*A, *Pcb*1692Δ*exp*I*-*p*exp*I, *Pcb1*692Δ*car*C*-*p*car*C, *Pcb1*692Δ*pyo*-p*pyo, Pcb1*692Δ*pyo*I-p*pyo*I and *Pcb1*692ΔT6pT6. Plasmid pJET4-*car*FG was transformed into *D. dadantii* in order to experimentally determine if the CarF and CarG proteins confer immunity to the carbapenem produced by *Pcb*1692. All transformants were maintained in LB supplemented with 15 µg/ml tetracycline.

### *In planta* competition assay

*In planta* competition assays were performed as previously described (41) with slight modifications. Targeted bacteria (*D. chrysanthami, D. dadantii, Pcc, P. atrosepticum* and *Pcb*G4P5 were transformed with plasmid pMP7605 conferring gentamycin resistance (123), as previously described. Wild-type *Pcb*1692, *Pcb*1692ΔT6 and *Pcb*1692ΔT6pT6 strains were grown overnight in LB supplemented with appropriate antibiotics. The optical density of overnight bacterial cultures was adjusted to OD_600_ = 1, resuspended in 1X PBS, mixed in a 1:1 ratio and inoculated into surface sterilized potato tubers (cv. Mondial). Inoculated tubers were placed in moist plastic containers, sealed and incubated for 72 h at 25°C. Macerated tuber tissue was scooped out and the CFU/ml of surviving targeted bacteria determined by serial dilutions on LB supplemented with gentamycin (15µg/ml). The assays were performed three independent times each consisting of three technical repeats.

### Detection of carbapenem and bacteriocin production by *Pcb*1692

Carbapenem production was determined by spot-on-lawn assay. Carbapenem production was determined for the following strains: wild-type *Pcb*1692, *Pcb*1692 mutant strains, HPI01 and laboratory stocks of *Pcb* strains whose genome sequences have not been determined (strains CC1, CC2, XT3, XT10, 358, G4P5 and G4P7). In summary, 10µl of *Pcb* strains was spotted on a lawn of freshly prepared targeted bacteria and a clear zone around *Pcb* was indicative of carbapenem production. To determine the role of iron in carbapenem production, lawns of targeted bacteria were prepared in M9 or M9 supplemented with iron (10µM). *Pcb* strains were then spotted onto the M9 and production of a clear zone determined 24hrs post inoculation at 28°C. Bacteriocin production was determined by spot-on-lawn overlay method, as previously described (129). In summary, 10µl of overnight cultures of *Pcb* strains were spotted on LB or M9, air dried and incubated overnight. Thereafter, bacteriocin production was induced by either UV irradiation or mitomycin C treatment. For mitomycin C induction, 10µl of mitomycin C (50µg/ml) was inoculated on top of *Pcb* strains, air dried and incubated in the dark for 5hrs. For UV irradiation, *Pcb* strains were exposed to UV light for 10sec and incubated in the dark for 5hrs. Thereafter, bacteriocin-induced *Pcb* cultures were killed with chloroform fumes for 10 minutes and then air dried for 30 minutes. Twenty microliters of overnight cultures of targeted bacteria was inoculated into 25ml of cooled 0.7% agar and overlayed on bacteriocin induced *Pcb*1692 strains. Clear zones around *Pcb* strains were indicative of bacteriocin production. In control experiments, *Pcb* strains were not exposed to UV or mitomycin C. All experiments were repeated three times. To determine the ability of *Pcb*1692 strains to produce carbapenem and bacteriocins under anaerobic conditions, the experimental procedures were the same as above except that bacteria were grown anaerobically in anaerobic jar (Merk) containing moistened Anaerocult^®^ A (Merk) which absorbs oxygen and moistened blue Anaerotest^®^ strip (Merk) which turns white in the absence of oxygen. Anaerobic jars were then tightly sealed and incubated overnight.

### Statistical analysis

All experiments were performed in triplicates and three independent times. Where applicable, an unpaired, two-tailed Student’s *t*-test was performed to determine statistical significance and a *p*-value less than 0.05 (*p*<0.05) was considered to be a statistically significant difference.

## Supporting information

## Acknowledgments

This research study was funded by the National Research Foundation (NRF), South Africa through Competitive Funding for Rated Researchers (CFRR) 98993; Bioinformatics and Functional Genomics (BFG 93685), Foundational Biodiversity Information Programmes (FBIP 86951) and The Technology Transfer Fund (RTF) 98654. DYS NRF BFG Post-Doctoral Fellowship, DB-R University of Pretoria Post-Doctoral Fellowship. NN, ARG and NLS were jointly funded by Potato South Africa Transformation Bursary and the NRF Grant Holder Linked Bursary. Any opinion, findings, conclusions or recommendations expressed in this material is that of the author(s) and the NRF does not accept any liability in this regard.

## Author contributions

The study was conceived and experiments designed by DYS and LNM. NLS performed metagenomic study. DB-R analysed metagenomics data and performed the *in silico* analysis. DYS, NN and ARG performed the experiments. DB-R, and LNM analysed the data and provided technical and scientific discussion: DYS wrote the manuscript. NN, ARG, DB-R and LNM revised the manuscript. All authors read and approved the final manuscript.

---

**S1 table. Summary of orthologs of the carbapenem and S-type pyosin in different SREs.** Sheet-1 shows orthology relationship of the carbapenem gene cluster in several SRE. Identified functional domains found in each gene in the car gene cluster is indicated. Sheet-2 shows bacteria strains with the homologous S-type pyosin identified in *Pcb*1692.

**S2 table. Putative Fur binding sites.** *In silico* analysis to identify putative Fur binding associated with the *car*R, *exp*I, *exp*R and *sly*A gene homologs of the *Pectobacterium brasiliense*1692 (*Pcb*1692) (sheet1), *Pectobacterium betavasculorum* (*Pbet*) ncppb2795 Sheet 2), *Pectobacterium atrosepticum* (*Pca*) icmp19972 (Sheet 3) and *Pectobacterium carotovorum* subsp. *carotovorum* (*Pcc*) ncppb312. Possible genomic hits and the consensus sequences are provided in the table.

**S3 table. Host range of killing by the *Pcb*1692 bacteriocin.** *Pcb*1692 and its isogenic mutant strains were spotted on LB and grown overnight. Bacteriocin production was induced with mitomycin C for 5hrs. Induced cells kill with chloroform vapour and overlaid with different bacteria. Clear zones are indicative of bacteriocin production.

## References

1. Glazebrook J, Roby D. Plant biotic interactions: from conflict to collaboration. The Plant Journal. 2018;93(4):589–91.

2. Xin X-F, Kvitko B, He SY. Pseudomonas syringae: what it takes to be a pathogen. Nature Reviews Microbiology. 2018;16(5):316.

3. Rodriguez-Moreno L, Ebert MK, Bolton MD, Thomma BP. Tools of the crook-infection strategies of fungal plant pathogens. The Plant Journal. 2018;93(4):664–74.

4. Bhunia AK. Foodborne microbial pathogens: mechanisms and pathogenesis: Springer; 2018.

5. Clayton JB, Al-Ghalith GA, Long HT, Van Tuan B, Cabana F, Huang H, et al. Associations Between Nutrition, Gut Microbiome, and Health in A Novel Nonhuman Primate Model. Scientific reports. 2018;8.

6. Bost A, Martinson VG, Franzenburg S, Adair KL, Albasi A, Wells MT, et al. Functional variation in the gut microbiome of wild Drosophila populations. Molecular ecology. 2018.

7. Kõiv V, Roosaare M, Vedler E, Kivistik PA, Toppi K, Schryer DW, et al. Microbial population dynamics in response to Pectobacterium atrosepticum infection in potato tubers. Scientific reports. 2015;5:11606.

8. Knief C, Delmotte N, Chaffron S, Stark M, Innerebner G, Wassmann R, et al. Metaproteogenomic analysis of microbial communities in the phyllosphere and rhizosphere of rice. The ISME journal. 2012;6(7):1378.

9. Yu K, Zhang T. Metagenomic and metatranscriptomic analysis of microbial community structure and gene expression of activated sludge. PloS one. 2012;7(5):e38183.

10. García-Bayona L, Comstock LE. Bacterial antagonism in host-associated microbial communities. Science. 2018;361(6408):eaat2456.

11. Stubbendieck RM, Straight PD. Escape from lethal bacterial competition through coupled activation of antibiotic resistance and a mobilized subpopulation. PLoS genetics. 2015;11(12):e1005722.

12. Bauer MA, Kainz K, Carmona-Gutierrez D, Madeo F. Microbial wars: Competition in ecological niches and within the microbiome. Microbial Cell. 2018;5(5):215.

13. Tyc O, Song C, Dickschat JS, Vos M, Garbeva P. The ecological role of volatile and soluble secondary metabolites produced by soil bacteria. Trends in microbiology. 2017;25(4):280–92.

14. Cornforth DM, Foster KR. Competition sensing: the social side of bacterial stress responses. Nature Reviews Microbiology. 2013;11(4):285.

15. Martin M, Dragoš A, Hölscher T, Maróti G, Bálint B, Westermann M, et al. De novo evolved interference competition promotes the spread of biofilm defectors. Nature communications. 2017;8:15127.

16. Hibbing ME, Fuqua C, Parsek MR, Peterson SB. Bacterial competition: surviving and thriving in the microbial jungle. Nature Reviews Microbiology. 2010;8(1):15.

17. Russell AB, Hood RD, Bui NK, LeRoux M, Vollmer W, Mougous JD. Type VI secretion delivers bacteriolytic effectors to target cells. Nature. 2011;475(7356):343–7.

18. Trunk K, Peltier J, Liu Y-C, Dill BD, Walker L, Gow NA, et al. The type VI secretion system deploys antifungal effectors against microbial competitors. Nature microbiology. 2018;3(8):920–31.

19. Hachani A, Wood TE, Filloux A. Type VI secretion and anti-host effectors. Current opinion in microbiology. 2016;29:81-93.

20. Russell AB, Peterson SB, Mougous JD. Type VI secretion system effectors: poisons with a purpose. Nature Reviews Microbiology. 2014;12(2):137–48.

21. Costa TR, Felisberto-Rodrigues C, Meir A, Prevost MS, Redzej A, Trokter M, et al. Secretion systems in Gram-negative bacteria: structural and mechanistic insights. Nature Reviews Microbiology. 2015;13(6):343.

22. Aoki SK, Diner EJ, de Roodenbeke CtK, Burgess BR, Poole SJ, Braaten BA, et al. A widespread family of polymorphic contact-dependent toxin delivery systems in bacteria. Nature. 2010;468(7322):439.

23. Desriac F, Defer D, Bourgougnon N, Brillet B, Le Chevalier P, Fleury Y. Bacteriocin as weapons in the marine animal-associated bacteria warfare: inventory and potential applications as an aquaculture probiotic. Marine drugs. 2010;8(4):1153–77.

24. Grinter R, Milner J, Walker D. Ferredoxin containing bacteriocins suggest a novel mechanism of iron uptake in Pectobacterium spp. PloS one. 2012;7(3):e33033.

25. Shyntum DY, Theron J, Venter SN, Moleleki LN, Toth IK, Coutinho TA. Pantoea ananatis utilizes a type VI secretion system for pathogenesis and bacterial competition. Molecular Plant-Microbe Interactions. 2015;28(4):420–31.

26. Coulthurst SJ, Barnard AM, Salmond GP. Regulation and biosynthesis of carbapenem antibiotics in bacteria. Nature Reviews Microbiology. 2005;3(4):295.

27. Verster AJ, Ross BD, Radey MC, Bao Y, Goodman AL, Mougous JD, et al. The landscape of type VI secretion across human gut microbiomes reveals its role in community composition. Cell host & microbe. 2017;22(3):411-9. e4.

28. Bernal P, Llamas MA, Filloux A. Type VI secretion systems in plant-associated bacteria. Environmental Microbiology. 2018;20(1):1–15.

29. Wall D. Kin recognition in bacteria. Annual review of microbiology. 2016;70:143–60.

30. Lutgring JD, Limbago BM. The problem of carbapenemase producing carbapenem-resistant Enterobacteriaceae detection. Journal of clinical microbiology. 2016:JCM. 02771-15.

31. Joshi A, Kostiuk B, Rogers A, Teschler J, Pukatzki S, Yildiz FH. Rules of engagement: the type VI secretion system in Vibrio cholerae. Trends in microbiology. 2017;25(4):267–79.

32. Shanker E, Federle MJ. Quorum sensing regulation of competence and bacteriocins in Streptococcus pneumoniae and mutans. Genes. 2017;8(1):15.

33. Charkowski AO. The Changing Face of Bacterial Soft-Rot Diseases. Annual review of phytopathology. 2018;56:269–88.

34. Adeolu M, Alnajar S, Naushad S, Gupta RS. Genome-based phylogeny and taxonomy of the ‘Enterobacteriales’: proposal for Enterobacterales ord. nov. divided into the families Enterobacteriaceae, Erwiniaceae fam. nov., Pectobacteriaceae fam. nov., Yersiniaceae fam. nov., Hafniaceae fam. nov., Morganellaceae fam. nov., and Budviciaceae fam. nov. International journal of systematic and evolutionary microbiology. 2016;66(12):5575–99.

35. van der Merwe JJ, Coutinho TA, Korsten L, van der Waals JE. Pectobacterium carotovorum subsp. brasiliensis causing blackleg on potatoes in South Africa. European Journal of Plant Pathology. 2010;126(2):175–85.

36. Panda P, Fiers M, Armstrong K, Pitman A. First report of blackleg and soft rot of potato caused by Pectobacterium carotovorum subsp. brasiliensis in New Zealand. New Dis Rep. 2012;26(15):2044-0588.2012.

37. Moraes A, Souza E, Mariano R, Silva A, Lima N, Peixoto A, et al. First Report of Pectobacterium aroidearum and Pectobacterium carotovorum subsp. brasiliensis Causing Soft Rot of Cucurbita pepo in Brazil. Plant Disease. 2017;101(2):379.

38. Voronina MV, Kabanova AP, Shneider MM, Korzhenkov AA, Toschakov SV, Miroshnikov KK, et al. First Report of Pectobacterium carotovorum subsp. brasiliense Causing Blackleg and Stem Rot Disease of Potato in Russia. Plant Disease. 2018(ja).

39. Ma X, Schloop A, Swingle B, Perry KL. Pectobacterium and Dickeya Responsible for Potato Blackleg Disease in New York State in 2016. Plant disease. 2018;102(9):1834–40.

40. Onkendi EM, Moleleki LN. Characterization of Pectobacterium carotovorum subsp. carotovorum and brasiliense from diseased potatoes in Kenya. European journal of plant pathology. 2014;139(3):557–66.

41. Marquez-Villavicencio MdP, Groves RL, Charkowski AO. Soft rot disease severity is affected by potato physiology and Pectobacterium taxa. Plant Disease. 2011;95(3):232–41.

42. Durrant A. Antimicrobial production by Pectobacterium carotovorum subspecies brasiliensis and its role in competitive fitness of the potato pathogen: Lincoln University; 2016.

43. Quast C, Pruesse E, Yilmaz P, Gerken J, Schweer T, Yarza P, et al. The SILVA ribosomal RNA gene database project: improved data processing and web-based tools. Nucleic acids research. 2012;41(D1):D590-D6.

44. Luis G, Rubio C, González-Weller D, Gutiérrez AJ, Revert C, Hardisson A. Comparative study of the mineral composition of several varieties of potatoes (Solanum tuberosum L.) from different countries cultivated in Canary Islands (Spain). International journal of food science & technology. 2011;46(4):774–80.

45. Tack FM. Trace elements in potato. Potato Research. 2014;57(3-4):311-25.

46. Fones H, Preston GM. The impact of transition metals on bacterial plant disease. FEMS Microbiology Reviews. 2013;37(4):495–519.

47. Bellieny-Rabelo D, Tanui CK, Miguel N, Kwenda S, Shyntum DY, Moleleki LN. Transcriptome and comparative genomics analyses reveal new functional insights on key determinants of pathogenesis and interbacterial competition in Pectobacterium and Dickeya spp. Appl Environ Microbiol. 2018.

48. Bateman A, Coin L, Durbin R, Finn RD, Hollich V, Griffiths-Jones S, et al. The Pfam protein families database. Nucleic acids research. 2004;32(suppl_1):D138–D41.

49. Bailey TL, Williams N, Misleh C, Li WW. MEME: discovering and analyzing DNA and protein sequence motifs. Nucleic acids research. 2006;34(suppl_2):W369-W73.

50. Rivet MM. Investigation into the regulation of exoenzyme production in Erwinia carotovora subspecies carotovora: University of Warwick; 1998.

51. Hamed RB, Gomez-Castellanos JR, Henry L, Ducho C, McDonough MA, Schofield CJ. The enzymes of ß-lactam biosynthesis. Natural Product Reports. 2013;30(1):21–107.

52. Chan Y-C, Wu J-L, Wu H-P, Tzeng K-C, Chuang D-Y. Cloning, purification, and functional characterization of Carocin S2, a ribonuclease bacteriocin produced by Pectobacterium carotovorum. BMC microbiology. 2011;11(1):99.

53. Chuang D-y, Chien Y-c, Wu H-P. Cloning and expression of the Erwinia carotovora subsp. carotovora gene encoding the low-molecular-weight bacteriocin carocin S1. Journal of bacteriology. 2007;189(2):620–6.

54. Axelrood PE, Rella M, Schroth MN. Role of antibiosis in competition of Erwinia strains in potato infection courts. Applied and environmental microbiology. 1988;54(5):1222–9.

55. Chassaing B, Cascales E. Antibacterial weapons: targeted destruction in the microbiota. Trends in microbiology. 2018;26(4):329–38.

56. Stubbendieck RM, Straight PD. Multifaceted interfaces of bacterial competition. Journal of bacteriology. 2016;198(16):2145–55.

57. Oluoch KR, Okanya PW, Hatti-Kaul R, Mattiasson B, Mulaa FJ. Protease-, Pectinase-and Amylase-Producing Bacteria from a Kenyan Soda Lake. The Open Biotechnology Journal. 2018;12(1).

58. Sundarram A, Murthy TPK. α-amylase production and applications: a review. Journal of Applied & Environmental Microbiology. 2014;2(4):166–75.

59. Dobson A, Cotter PD, Ross RP, Hill C. Bacteriocin production: a probiotic trait? Appl Environ Microbiol. 2012;78(1):1–6.

60. Silverman JM, Brunet YR, Cascales E, Mougous JD. Structure and regulation of the type VI secretion system. Annual review of microbiology. 2012;66:453-72.

61. Ishikawa T, Sabharwal D, Bröms J, Milton DL, Sjöstedt A, Uhlin BE, et al. Pathoadaptive conditional regulation of the type VI secretion system in Vibrio cholerae O1 strains. Infection and immunity. 2012;80(2):575–84.

62. Gueguen E, Durand E, Zhang XY, d’Amalric Q, Journet L, Cascales E. Expression of a Yersinia pseudotuberculosis type VI secretion system is responsive to envelope stresses through the OmpR transcriptional activator. PloS one. 2013;8(6):e66615.

63. Si M, Zhao C, Burkinshaw B, Zhang B, Wei D, Wang Y, et al. Manganese scavenging and oxidative stress response mediated by type VI secretion system in Burkholderia thailandensis. Proceedings of the National Academy of Sciences. 2017:201614902.

64. Chakraborty S, Sivaraman J, Leung KY, Mok YK. The two-component PhoB-PhoR regulatory system and ferric uptake regulator sense phosphate and iron to control virulence genes in type III and VI secretion systems of Edwardsiella tarda. Journal of Biological Chemistry. 2011:jbc. M111. 295188.

65. Leonard S, Hommais F, Nasser W, Reverchon S. Plant–phytopathogen interactions: bacterial responses to environmental and plant stimuli. Environmental microbiology. 2017;19(5):1689–716.

66. Erill I, Campoy S, Barbé J. Aeons of distress: an evolutionary perspective on the bacterial SOS response. FEMS microbiology reviews. 2007;31(6):637–56.

67. Thorgersen MP, Lancaster WA, Ge X, Zane GM, Wetmore KM, Vaccaro BJ, et al. Mechanisms of Chromium and Uranium Toxicity in Pseudomonas stutzeri RCH2 Grown under Anaerobic Nitrate-Reducing Conditions. Frontiers in Microbiology. 2017;8:1529.

68. Leaden L, Silva LG, Ribeiro RA, dos Santos NM, Lorenzetti AP, Alegria TG, et al. Iron Deficiency Generates Oxidative Stress and Activation of the SOS Response in Caulobacter crescentus. Frontiers in microbiology. 2018;9.

69. Fang Y, Mercer RG, McMullen LM, Gänzle MG. Induction of Shiga toxin prophage by abiotic environmental stress in food. Applied and environmental microbiology. 2017:AEM. 01378-17.

70. Hugouvieux-Cotte-Pattat N, Condemine G, Shevchik VE. Bacterial pectate lyases, structural and functional diversity. Environmental microbiology reports. 2014;6(5):427–40.

71. Effantin G, Rivasseau C, Gromova M, Bligny R, Hugouvieux-Cotte-Pattat N. Massive production of butanediol during plant infection by phytopathogenic bacteria of the genera Dickeya and Pectobacterium. Molecular microbiology. 2011;82(4):988–97.

72. Pi H, Helmann JD. Ferrous iron efflux systems in bacteria. Metallomics. 2017;9(7):840–51.

73. Wierzbowska J, Rychcik B, Światly A. The effect of different production systems on the content of micronutrients and trace elements in potato tubers. Acta Agriculturae Scandinavica, Section B—Soil & Plant Science. 2018:1-8.

74. Cooper B, Islam N, Xu Y, Beard HS, Garrett WM, Gu G, et al. Quantitative Proteomic Analysis of Staphylococcus aureus Treated With Punicalagin, a Natural Antibiotic From Pomegranate That Disrupts Iron Homeostasis and Induces SOS. Proteomics. 2018;18(9):1700461.

75. Goyal RK, Mattoo AK. Multitasking antimicrobial peptides in plant development and host defense against biotic/abiotic stress. Plant Science. 2014;228:135-49.

76. McGowan SJ, Barnard AM, Bosgelmez G, Sebaihia M, Simpson NJ, Thomson NR, et al. Carbapenem antibiotic biosynthesis in Erwinia carotovora is regulated by physiological and genetic factors modulating the quorum sensing-dependent control pathway. Molecular microbiology. 2005;55(2):526–45.

77. McGowan S, Sebaihia M, Jones S, Yu B, Bainton N, Chan P, et al. Carbapenem antibiotic production in Erwinia carotovora is regulated by CarR, a homologue of the LuxR transcriptional activator. Microbiology. 1995;141(3):541–50.

78. Lee DH, Kim J-B, Lim J-A, Han S-W, Heu S. Genetic diversity of Pectobacterium carotovorum subsp. brasiliensis isolated in Korea. The Plant pathology journal. 2014;30(2):117.

79. Andrews SC, Robinson AK, Rodríguez-Quiñones F. Bacterial iron homeostasis. FEMS microbiology reviews. 2003;27(2-3):215-37.

80. Seo SW, Kim D, Latif H, O’Brien EJ, Szubin R, Palsson BO. Deciphering Fur transcriptional regulatory network highlights its complex role beyond iron metabolism in Escherichia coli. Nature communications. 2014;5:4910.

81. Niehus R, Picot A, Oliveira NM, Mitri S, Foster KR. The evolution of siderophore production as a competitive trait. Evolution. 2017;71(6):1443–55.

82. Chandrangsu P, Rensing C, Helmann JD. Metal homeostasis and resistance in bacteria. Nature Reviews Microbiology. 2017;15(6):338.

83. Foster JW. Salmonella acid shock proteins are required for the adaptive acid tolerance response. Journal of Bacteriology. 1991;173(21):6896–902.

84. Hassan HM, Troxell B. Transcriptional regulation by Ferric Uptake Regulator (Fur) in pathogenic bacteria. Frontiers in cellular and infection microbiology. 2013;3:59.

85. Porcheron G, Dozois CM. Interplay between iron homeostasis and virulence: Fur and RyhB as major regulators of bacterial pathogenicity. Veterinary microbiology. 2015;179(1-2):2–14.

86. Delany I, Rappuoli R, Scarlato V. Fur functions as an activator and as a repressor of putative virulence genes in Neisseria meningitidis. Molecular microbiology. 2004;52(4):1081–90.

87. Massé E, Vanderpool CK, Gottesman S. Effect of RyhB small RNA on global iron use in Escherichia coli. Journal of bacteriology. 2005;187(20):6962–71.

88. Carpenter BM, Whitmire JM, Merrell DS. This is not your mother’s repressor: the complex role of fur in pathogenesis. Infection and immunity. 2009;77(7):2590–601.

89. Embree M, Qiu Y, Shieu W, Nagarajan H, O’Neil R, Lovley D, et al. The iron stimulon and fur regulon of Geobacter sulfurreducens and their role in energy metabolism. Applied and environmental microbiology. 2014:AEM. 03916-13.

90. Skaar EP. The battle for iron between bacterial pathogens and their vertebrate hosts. PLoS pathogens. 2010;6(8):e1000949.

91. Dunham NP, Chang W-c, Mitchell AJ, Martinie RJ, Zhang B, Bergman JA, et al. Two Distinct Mechanisms for C–C Desaturation by Iron (II)-and 2-(Oxo) glutarate-Dependent Oxygenases: Importance of a-Heteroatom Assistance. Journal of the American Chemical Society. 2018;140(23):7116–26.

92. Massé E, Vanderpool CK, Gottesman S. Effect of RyhB Small RNA on Global Iron Use in Escherichia coli. Journal of Bacteriology. 2005;187(20):6962–71.

93. Cui Y, Chatterjee A, Hasegawa H, Dixit V, Leigh N, Chatterjee AK. ExpR, a LuxR homolog of Erwinia carotovora subsp. carotovora, activates transcription of rsmA, which specifies a global regulatory RNA-binding protein. Journal of bacteriology. 2005;187(14):4792–803.

94. Mole BM, Baltrus DA, Dangl JL, Grant SR. Global virulence regulation networks in phytopathogenic bacteria. Trends in microbiology. 2007;15(8):363–71.

95. Cui Y, Chatterjee A, Liu Y, Dumenyo CK, Chatterjee AK. Identification of a global repressor gene, rsmA, of Erwinia carotovora subsp. carotovora that controls extracellular enzymes, N-(3-oxohexanoyl)-L-homoserine lactone, and pathogenicity in soft-rotting Erwinia spp. Journal of bacteriology. 1995;177(17):5108–15.

96. Wilf NM, Reid AJ, Ramsay JP, Williamson NR, Croucher NJ, Gatto L, et al. RNA-seq reveals the RNA binding proteins, Hfq and RsmA, play various roles in virulence, antibiotic production and genomic flux in Serratia sp. ATCC 39006. BMC genomics. 2013;14(1):1.

97. Andersson RA, Eriksson AR, Heikinheimo R, Mäe A, Pirhonen M, Kõiv V, et al. Quorum sensing in the plant pathogen Erwinia carotovora subsp. carotovora: the role of expREcc. Molecular plant-microbe interactions. 2000;13(4):384–93.

98. Miller J. Miller JH.. Experiments in Molecular Genetics. Cold Spring Harbor Laboratory Press, Cold Spring Harbor, NY; 1972.

99. Duarte V, de Boer SH, Ward LJ, de Oliveira AM. Characterization of atypical Erwinia carotovora strains causing blackleg of potato in Brazil. J Appl Microbiol. 2004;96(3):535–45.

100. Onkendi EM, Ramesh AM, Kwenda S, Naidoo S, Moleleki L. Draft Genome Sequence of a Virulent Pectobacterium carotovorum subsp. brasiliense Isolate Causing Soft Rot of Cucumber. Genome Announc. 2016;4(1).

101. .!!! INVALID CITATION !!! {}.

102. Gardan L, Gouy C, Christen R, Samson Rg. Elevation of three subspecies of Pectobacterium carotovorum to species level: Pectobacterium atrosepticum sp. nov., Pectobacterium betavasculorum sp. nov. and Pectobacterium wasabiae sp. nov. International journal of systematic and evolutionary microbiology. 2003;53(2):381–91.

103. Goto M, Matsumoto K. Erwinia carotovora subsp. wasabiae subsp. nov. isolated from diseased rhizomes and fibrous roots of Japanese horseradish (Eutrema wasabi Maxim.). International Journal of Systematic and Evolutionary Microbiology. 1987;37(2):130–5.

104. Hori S. A bacterial leaf-disease of tropical orchids. Zentralblatt für Bakteriologie, Parasitenkunde und Infektionskrankheiten. 1911;31:85-92.

105. Hauben L, Moore ER, Vauterin L, Steenackers M, Mergaert J, Verdonck L, et al. Phylogenetic position of phytopathogens within the Enterobacteriaceae. Systematic and Applied Microbiology. 1998;21(3):384–97.

106. Samson R, Legendre J, Christen R, Fischer-Le Saux M, Achouak W, Gardan L. 1973 and Brenneria paradisiaca to the genus Dickeya gen. nov. as Dickeya chrysanthemi comb. nov. and Dickeya paradisiaca comb. nov. and delineation of four novel species, Dickeya dadantii sp. nov., Dickeya dianthicola sp. nov., Dickeya dieffenbachiae sp. nov. and Dickeya zeae sp. nov. International Journal of Systematic and Evolutionary Microbiology. 2005;55:1415–27.

107. Burkholder W, McFadden LA, Dimock E. A bacterial blight ol Chrysanthemums. Phytopathology. 1953;43(9).

108. Gibbons R, Engle L. Vitamin K in Aerobacter aerogenes and Serratia marcescens. Journal of bacteriology. 1965;90(2):561.

109. Grimont PA, Grimont F, Starr MP. Serratia species isolated from plants. Current Microbiology. 1981;5(5):317–22.

110. Serrano F. Bacterial fruitlet brown-rot of Pineapple in the Philippines. Philippine Journal of Science. 1928;36(3).

111. Popp A, Cleenwerck I, Iversen C, De Vos P, Stephan R. Pantoea gaviniae sp. nov. and Pantoea calida sp. nov., isolated from infant formula and an infant formula production environment. International journal of systematic and evolutionary microbiology. 2010;60(12):2786–92.

112. Fadrosh DW, Ma B, Gajer P, Sengamalay N, Ott S, Brotman RM, et al. An improved dual-indexing approach for multiplexed 16S rRNA gene sequencing on the Illumina MiSeq platform. Microbiome. 2014;2(1):6.

113. Klindworth A, Pruesse E, Schweer T, Peplies J, Quast C, Horn M, et al. Evaluation of general 16S ribosomal RNA gene PCR primers for classical and next-generation sequencing-based diversity studies. Nucleic acids research. 2013;41(1):e1–e.

114. Bolger AM, Lohse M, Usadel B. Trimmomatic: a flexible trimmer for Illumina sequence data. Bioinformatics. 2014;30(15):2114–20.

115. Martin M. Cutadapt removes adapter sequences from high-throughput sequencing reads. EMBnetjournal, North America. 17. 2011.

116. Camacho C, Coulouris G, Avagyan V, Ma N, Papadopoulos J, Bealer K, et al. BLAST+: architecture and applications. BMC bioinformatics. 2009;10(1):421.

117. Ondov BD, Bergman NH, Phillippy AM. Interactive metagenomic visualization in a Web browser. BMC bioinformatics. 2011;12(1):385.

118. Mattinen L, Nissinen R, Riipi T, Kalkkinen N, Pirhonen M. Host-extract induced changes in the secretome of the plant pathogenic bacterium Pectobacterium atrosepticum. Proteomics. 2007;7(19):3527–37.

119. Li L, Stoeckert CJ, Jr., Roos DS. OrthoMCL: identification of ortholog groups for eukaryotic genomes. Genome Res. 2003;13(9):2178–89.

120. Munch R, Hiller K, Barg H, Heldt D, Linz S, Wingender E, et al. PRODORIC: prokaryotic database of gene regulation. Nucleic Acids Res. 2003;31(1):266–9.

121. Kazakov AE, Cipriano MJ, Novichkov PS, Minovitsky S, Vinogradov DV, Arkin A, et al. RegTransBase—a database of regulatory sequences and interactions in a wide range of prokaryotic genomes. Nucleic acids research. 2006;35(suppl_1):D407-D12.

122. Kilic S, White ER, Sagitova DM, Cornish JP, Erill I. CollecTF: a database of experimentally validated transcription factor-binding sites in Bacteria. Nucleic Acids Res. 2014;42(Database issue):D156–60.

123. Lagendijk EL, Validov S, Lamers GE, De Weert S, Bloemberg GV. Genetic tools for tagging Gram-negative bacteria with mCherry for visualization in vitro and in natural habitats, biofilm and pathogenicity studies. FEMS microbiology letters. 2010;305(1):81–90.

124. Datsenko KA, Wanner BL. One-step inactivation of chromosomal genes in Escherichia coli K-12 using PCR products. Proceedings of the National Academy of Sciences. 2000;97(12):6640–5.

125. Tanui CK, Shyntum DY, Priem SL, Theron J, Moleleki LN. Influence of the ferric uptake regulator (Fur) protein on pathogenicity in Pectobacterium carotovorum subsp. brasiliense. PloS one. 2017;12(5):e0177647.

126. Moleleki LN, Pretorius RG, Tanui CK, Mosina G, Theron J. A quorum sensing-defective mutant of Pectobacterium carotovorum ssp. brasiliense 1692 is attenuated in virulence and unable to occlude xylem tissue of susceptible potato plant stems. Mol Plant Pathol. 2017;18(1):32–44.

127. Katashkina JI, Hara Y, Golubeva LI, Andreeva IG, Kuvaeva TM, Mashko SV. Use of the λ Red-recombineering method for genetic engineering of Pantoea ananatis. BMC molecular biology. 2009;10(1):34.

128. Heeb S, Itoh Y, Nishijyo T, Schnider U, Keel C, Wade J, et al. Small, stable shuttle vectors based on the minimal pVS1 replicon for use in gram-negative, plant-associated bacteria. Molecular Plant-Microbe Interactions. 2000;13(2):232–7.

129. Roh E, Lee S, Lee Y, Ra D, Choi J, Moon E, et al. Diverse antibacterial activity of Pectobacterium carotovorum subsp. carotovorum isolated in Korea. J Microbiol Biotechnol. 2009;19(1):42–50.

